# Temporal order of activations and interactions during arithmetic calculations measured by intracranial electrophysiological recordings in the human brain

**DOI:** 10.64898/2026.01.07.698095

**Authors:** M Kalinova, B Kerkova, A Kalina, V Pytelova, J Amlerova, R Janca, P Jezdik, D Krysl, M Kudr, P Krsek, P Marusic, J Hammer

**Author notes:** joint last authors (P Marusic, J Hammer).

## Abstract

Arithmetic requires complex and fast processes orchestrated within a large-scale network spanning multiple brain regions. However, reports on the network’s temporal dynamics are scarce. Here, we present data from intracranial EEG (iEEG) of 20 subjects (epilepsy surgery candidates) performing a sequential three-operand arithmetic task. Utilizing the high temporal and spatial resolution of iEEG, we analysed changes in high-gamma band (HGB; 52–120 Hz) activity and functional connectivity assessed by phase-locking value (PLV) in the delta (0.1–3 Hz) and theta (3–7 Hz) frequency bands. Strong and transient HGB activations peaked first in the ventral occipito-temporal cortex, followed by a more gradual increase in the lateral parietal, sensorimotor, and frontal cortices, accompanied by deactivations in default mode network areas. The connectivity patterns were more extensive during calculation than number recognition, with the theta PLV peaking ∼150 ms earlier than the delta PLV. Earliest connectivity appeared, surprisingly, between ventral temporal and frontal regions at ∼100–200 ms, evolving into a robust pattern among key network nodes at ∼200–400 ms after the presentation of each operand. The presented results elucidate information flow within the putative arithmetic network during calculation in the human brain, offering high-temporal-resolution insights into its functional architecture.

## Introduction

Arithmetic relies on a set of cognitive functions, which include basic numerical (recognizing numbers from symbols, decoding their value), and arithmetic abilities (operation recognition, fact retrieval), alongside more general cognitive skills such as working memory and decision making^1^. Thanks to extensive neuroimaging research, we know that regions supporting these processes form a widespread network in the human brain^2^, hereafter referred to as the “arithmetic network”. For clarity, the term “arithmetic network” is used as a conceptual simplification to denote the set of brain regions commonly implicated in arithmetic processing, consistent with prior usage^3^. This term overlaps with related concepts such as the “math network”^3^ or “math/arithmetic processing network”^4,5^. The regions encompassed by this network are unlikely to be exclusively dedicated to arithmetic processing; rather, they also contribute to, and overlap with, other cognitive functions and processes. Thus, the “arithmetic network” is considered here a functional network that emerges from dynamic interactions among distributed regions during arithmetic tasks.

Most research on the arithmetic network to date has utilized techniques like functional MRI (fMRI), which produce rather static activation maps based on slow changes in brain metabolism. However, the limited temporal resolution of these methods is likely to obscure critical dynamic network aspects of a process as fast and complex as arithmetic equation solving. Reports on the precise temporal dynamics of activations and connectivity patterns during such tasks remain scarce. Thus, an emphasis on the network dynamics, in terms of both activations and connectivity, is necessary for greater insight into the neural implementation of arithmetic.

The arithmetic network presumably consists of several distributed regions or network nodes (Fig. 1). These regions–each with a distinct and not necessarily arithmetic-specific role–interact to secure flexible and efficient numerical and arithmetic processing. Visual information is processed first in the ventral occipito-temporal cortex (VOTC)^6^, where highly specific activations for numerical symbols have been found in the posterior part of the inferior temporal gyrus (pITG), including the putative number form area. This area typically responds more to Arabic digits than other meaningful symbols like letters or semantically similar stimuli like number words^4,5^, although some suggest that this effect may also be explained by intervening factors like computational demands^7^.

**Figure 1:**
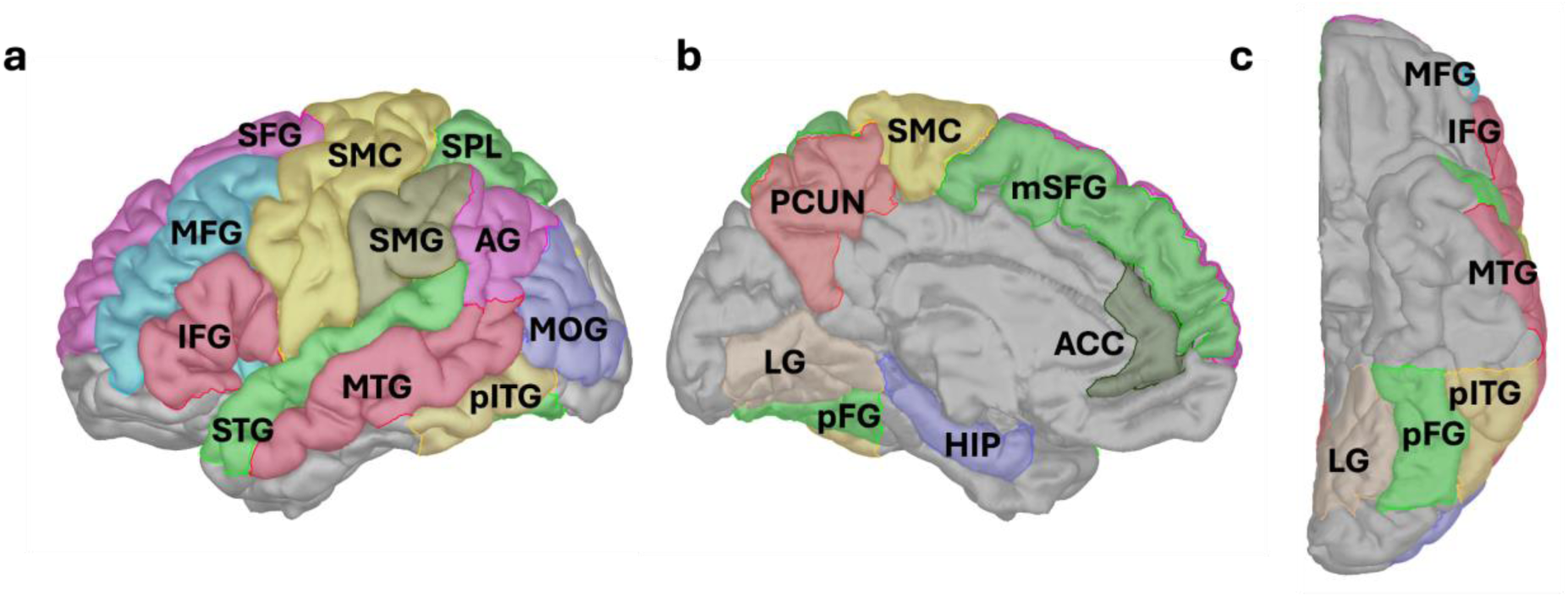
Regions of the putative arithmetic network. We selected 18 regions of interest (ROIs) associated with arithmetic processing, here depicted on a brain model (Colin27). **(a)** Lateral view of left hemisphere, **(b)** medial view, **(c)** bottom view, Abbreviations: MOG, middle occipital gyrus; pFG, posterior part of fusiform gyrus; LG, lingual gyrus; pITG, posterior part of inferior temporal gyrus; MTG, middle temporal gyrus; STG, superior temporal gyrus; IPS, intraparietal sulcus; SMG, supramarginal gyrus; AG, angular gyrus; PCUN, precuneus; SPL, superior parietal lobule; ACC, anterior cingulate cortex; SMC, sensorimotor cortex (precentral/postcentral gyri and central operculum); HIP, hippocampus; SFG, superior frontal gyrus; mSFG, medial part of superior frontal gyrus; MFG, middle frontal gyrus; IFG, inferior frontal gyrus. Notably, the arithmetic network is not unique to calculation, but also includes other domain-general processes that are closely linked to arithmetic problem solving (e.g., working memory, attention allocation).

Involvement of other VOTC regions, including the posterior part of the fusiform gyrus (pFG) and the lingual gyrus (LG), has also been reported^8,9^. Verbal number processing may be primarily subserved by the inferior parietal lobule, consisting of the angular gyrus (AG) and supramarginal gyrus (SMG)^10,11^. The intraparietal sulci (IPS) can be described as the “supramodal quantity hub” for numerical cognition, responsible for processing magnitudes and generating their semantic representations^11,12^. While the IPS is typically believed to hold purely abstract number representations, signs of format-dependency are occasionally also reported^10,13,14^.

Once the numbers are recognized, arithmetic calculations can be initiated. This yields widespread fronto-parietal engagement, indicative of the range of operations and associated cognitive strategies that calculations can encompass^9^. Some operations are presumably solved through direct retrieval from long-term memory; simple multiplication and single-digit addition are frequently ascribed to this strategy(^15,16^, but see also^17,18^). Retrieval-based problems primarily recruit the inferior parietal lobule, with additional involvement of the middle and superior temporal gyri (MTG, STG) and the hippocampus (HIP)^8,10,19–21^. In contrast, operations like subtraction, as well as more difficult problems like those involving larger operands or cross-decade results, depend more on procedural strategies, placing further demands on domain-general cognitive functions like working memory or cognitive control^15,22–25^. Correspondingly, procedural problems preferentially recruit frontal regions, including the inferior, middle, and superior frontal gyri (IFG, MFG, SFG, respectively)^1,26^, possibly also the anterior cingulate cortex (ACC)^9,20,27^. Greater or more widespread parietal responses, including in the superior parietal lobule (SPL) and precuneus (PCUN), can also reflect increasing problem difficulty or visuospatial attention demands^1,8,16,28,29^. As apparent, activity across the arithmetic network may vary with both problem characteristics and the domain-general cognitive demands they impose.

The sequence, duration, or temporal profile of activations in the different regions of the arithmetic network may help to specify the stages of these underlying cognitive processes. EEG shows that early components of event-related potentials are likely associated with the visual processing of digits, whereas later parietal and frontal activations correspond more closely with calculations, then scaling with problem difficulty^30–32^. Likewise, evoked activity measured by MEG during sequential arithmetic tasks showed an increase of global field potential after every stimulus, propagating from posterior to anterior regions^33^.

More detailed insight can be obtained from human intracranial EEG (iEEG) studies, which show that distinct regions of the arithmetic network differ in terms of the timing, magnitude and duration of high-gamma band (HGB) signals^34^. For example, neural activations in pITG were stronger following the second operand in a sequential arithmetic task, suggesting the involvement of this region in the manipulation of numbers rather than their mere recognition^35^. Daitch et al. disentangled math-selective sites within the pITG and anterior IPS and suggested their mutual functional coupling^6^. Recently, Pinheiro-Chagas et al. described successive activations in nine regions of the arithmetic network, showing a consistent temporal order of activations from VOTC to lateral parietal regions, and finally in frontal regions^3^.

Fast and flexible interactions among the regions of the arithmetic network are thought to be necessary for successful calculation^21^ and can be assessed by means of functional connectivity (FC). Results from fMRI again confirm a widespread network, including the parietal and frontal cortex^36,37^. These regions also interact with the hippocampus (HIP), which is important for memory-based arithmetic fact retrieval and for arithmetic skill acquisition (see^38^ for review). From the iEEG research, which is well-positioned to describe the fast temporal changes of FC, Daitch et al. reported the existence of bidirectional feedback loops between lateral parietal and ventral temporal cortical areas^6^. Das and Menon identified a network of several lateral parietal regions (IPS, SPL, and SMG) as a dominant inflow hub, with causal influences from the VOTC (especially FG) and HIP^21^. In the temporal sequence reported by Pinheiro-Chagas et al., higher cross-correlations were observed in temporally adjacent regions than those further apart in their sequential order^3^.

Despite these valuable reports, several key aspects of the arithmetic network’s dynamics are currently not well described. For example, it remains unclear whether there is an interpretable relationship between temporal activation profiles, the precise temporal sequence of activations and the roles of the different regions in the network. Similarly, the issue of numerical format dependence across the arithmetic network remains a matter of debate. Based on previous reports^5,10,13,39^, we hypothesized that the pITG will show higher specificity for Arabic numbers, whereas the brain activity in the IPS will be format independent. Regarding the FC within the arithmetic network, the evolution of connectivity patterns as well as the frequency bands utilized for long-range communication have yet to be identified. It is also unknown to which extent these patterns differ according to task, such as between recognizing the numerical information and calculation.

To capture the fast (i.e., millisecond-scale) dynamics of neuronal activations and connectivity changes during arithmetic computations, we analyzed iEEG signals from 20 subjects (epilepsy surgery candidates). A remarkable feature of the iEEG signal is the relatively robust estimate of HGB activity (here defined as iEEG power modulation between 52–120 Hz). While the exact mechanisms generating HGB remain a matter of debate, their most notable property has become well-established: they provide high temporal-resolution evidence for the participation of local neural populations during the task-at-hand^40,41^. The role of the HGB as a proxy for local neural activity is further supported by the band’s correlation and close anatomical correspondence with fMRI BOLD signals^42^. Moreover, the iEEG enables the simultaneous measurement of local neuronal population activity in distant brain areas, and the study of their FC changes in time.

To describe neural responses across the arithmetic network, we developed a sequential calculation task with three-operand equations. Since the operands (and operators) were presented sequentially, the presentation of the first operand was associated only with number recognition, while the other two operands required additional calculation. We included a range of numerical formats, which the subjects manipulated at varying levels of problem difficulty across two operation types: addition and subtraction. This has allowed us to test these additional effects, which may help characterize the functional contributions of individual regions.

Below, we describe different neural activity profiles of regions in the arithmetic network, the temporal order of their HGB activations, and temporal changes of FC patterns.

## 2 Results

### 2.1 Behavioural results

Twenty subjects (epilepsy surgery candidates; median age 36; range 18–63; see Suppl. Table T1 for further details) implanted with iEEG multicontact electrodes localized across the putative arithmetic network (Suppl. Figs. S1 and S2) performed the sequential arithmetic task (see Fig. 2 and Methods for more details). The sequentially presented arithmetic equations (e.g., 8 - 3 + 2 = 6: yes/no?) consisted of three operands and two operators (plus, minus). Mean reaction time was 1.0 ± 0.4 s (mean ± SD across subjects, range 0.6–2.1 s), and the ratio of the correct answers was 83 ± 3 % (mean ± SEM across subjects, range 47–96 %). In the analysis below, we focused on the arithmetic-related neural activity only up to the ‘=’ sign; the later response was excluded as it involved other confounding cognitive processes (e.g., results comparison, planning of motor actions (button press), movement execution, error monitoring, and feedback).

**Figure 2:**
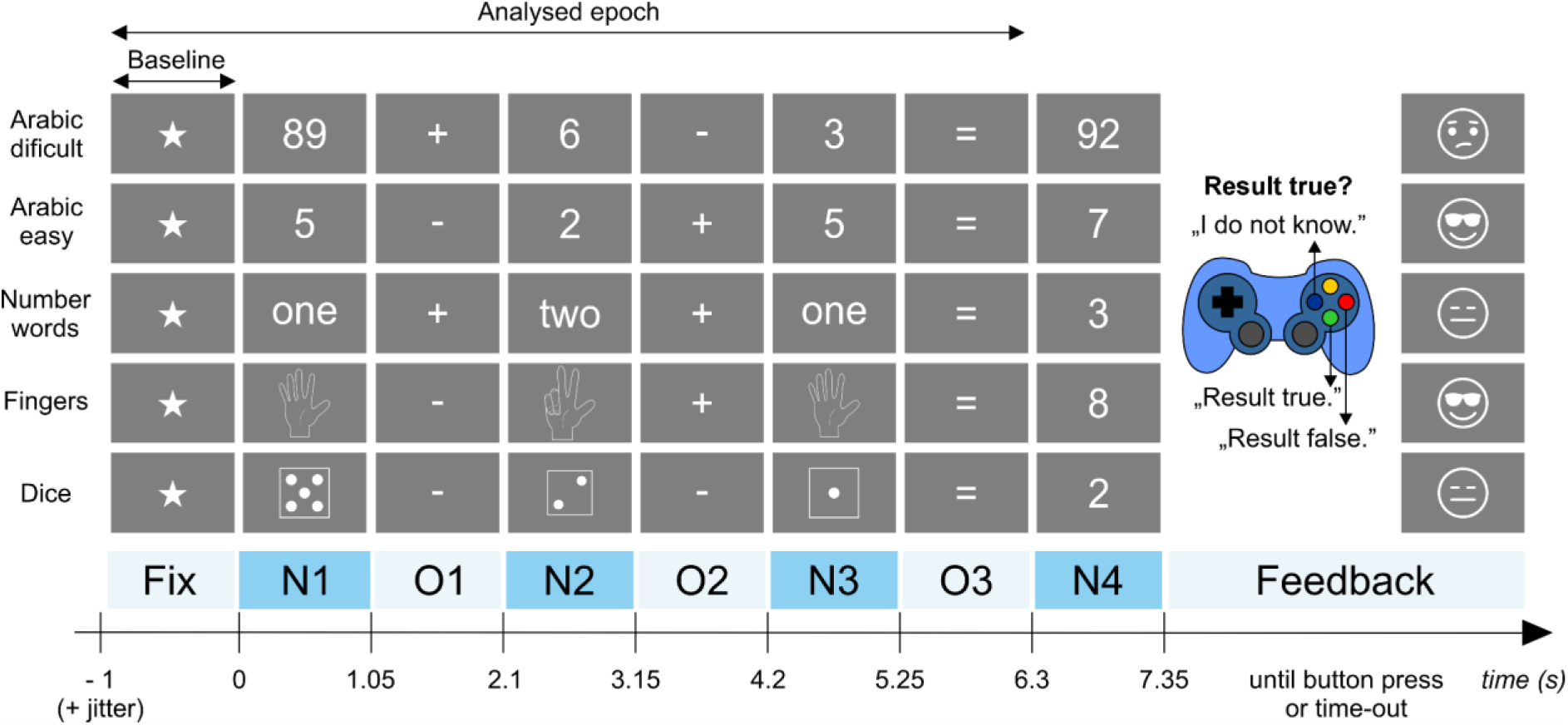
Sequential arithmetic task. The first five rows represent examples of the task, where trials consisted of sequentially presented equations with the same general structure (bottom row). First, a fixation cross (a star) appeared for 1.0–1.5 s (randomly varied), followed by the first operand - number (N1), first operator (O1), second operand (N2), second operator (O2), third operand (N3), equals sign (O3), and a proposed result (N4). Subjects were asked to answer whether the result was true or false by pressing a corresponding coloured button on a gamepad (red = false result; green = true result; blue = uncertain). Feedback was provided based on their answer. Equations were either difficult (N1 was a double digit, result was over ten, only Arabic numerals; presented on the top row) or easy (single Arabic digits, words or symbols, from 0–9, result never exceeded ten).

We restricted the analysis to correct trials only (N_trials_ = 96 ± 4, mean ± SEM across subjects). Alongside Arabic numerals, we included different numerical formats (number words, dice symbols and fingers), to address the issue if and to which extent the arithmetic-related representations are abstract or format-dependent in the human brain^26^. We found no statistical differences in the reaction times nor in the ratios of correct answers among the numerical formats for the easier arithmetic calculations (i.e., result < 10), but—as could be anticipated—the subjects were significantly slower and less accurate in the “Arabic difficult” condition, where the intermediate results were over a decade (see Suppl. Fig. S3).

### 2.2 Arithmetic network activity: high-gamma band temporal profiles

We selected 18 non-overlapping anatomical regions of interest (ROIs) of the human brain (based on literature reports; see Methods for details), presumed to form the major nodes of the distributed and large-scale arithmetic network (Fig. 1). Network activity was sampled by a total of 1,518 iEEG channels (i.e., bipolar-referenced iEEG electrode contacts). ROI-specific activations (spectrograms or HGB activity) were computed as the grand average across all trial-averaged activities of iEEG channels assigned to that ROI (see Suppl. Fig S2 and Methods for more details). Note that in order to uncover the widespread activation in the human brain, we compared activity (and FC) against a baseline period (i.e., the presentation of the fixation cross, similar to other, recent studies^3,43^, rather than using the subtractive approach of a control task (see Discussion).

First, we investigated the entire spectrograms of ROIs during the sequential arithmetic task (Fig. 3), regardless of the numerical format or operator type (see Sections 2.4). Increases in HGB power were observed, typically accompanied by a simultaneous decrease in power < 50 Hz, most pronounced in the alpha (8–12 Hz) and beta (13–30 Hz) bands. The opposite pattern was observed for HGB decreases, with low-frequency (< 13 Hz) power increases, most pronounced in the theta (3–7 Hz) band of the ACC. These patterns represent typical spectral fingerprints of cortical activation and deactivation^44^. However, there were also notable exceptions, for example in the STG. These results could also be of relevance to research using scalp EEG or MEG spectral power, where the HGB activity is generally harder to reliably estimate during cognitive tasks.

**Figure 3:**
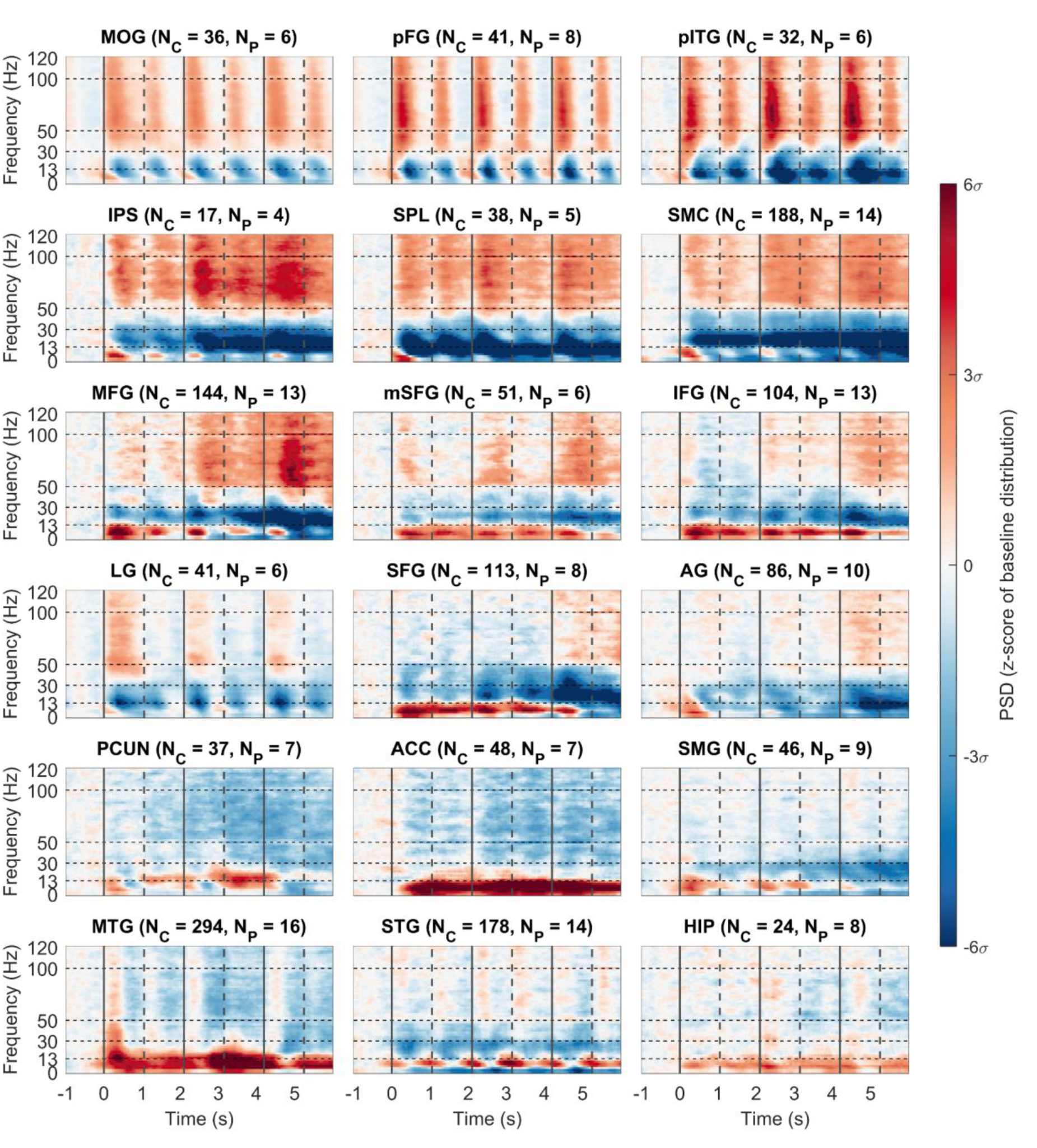
Spectrograms of arithmetic network ROIs during the sequential calculation task. Each subplot represents one ROI, indicated by the title with the number of channels and subjects used for the grand-average. The channel-averaged spectrogram of power spectrum density (PSD, colour-coded) in the time (x-axis) and frequency (y-axis) domains is displayed. The colour-limits of the PSD are expressed in the z-score of the baseline data distribution of each ROI separately. The baseline period was taken in the time interval from -1 s to -0.25 s. Vertical lines in spectrograms represent the presentation of the stimuli (solid line for operands and dashed lines for operators) of the sequential calculation task, with the first operator appearing at time t = 0 s). Abbreviations: N_c_, total number of channels in the ROI; N_P_, number of subjects contributing the channels to the ROI.

Next, we focused on the activity in the HGB (52–120 Hz), in the form of time-resolved power modulations. We observed several differential patterns of HGB activity across the arithmetic network ROIs (Fig. 4): (1) there were short, transient increases following the stimulus presentation with a rapid return to baseline values in the MOG and VOTC regions (LG, pITG, and pFG). The HGB peaks were higher after the presentation of operands than operators, consistent with other iEEG reports^3,6,35^. (2) More gradual increases were observed in IPS, SPL, and sensorimotor regions, as were (3) increases, most pronounced after the presentation of the third operand, in the frontal regions (mSFG, MFG, and IFG). Apart from activations, we also observed significant decreases in power in the PCUN and ACC. These network activity results were also corroborated at the level of trial-averaged, single-channel analysis (Suppl. Fig. S5), which showed that most activated network areas were the IPS, followed by the pITG, pFG, MOG, and SPL; this confirms their role as the “core” nodes of the arithmetic network.

**Figure 4:**
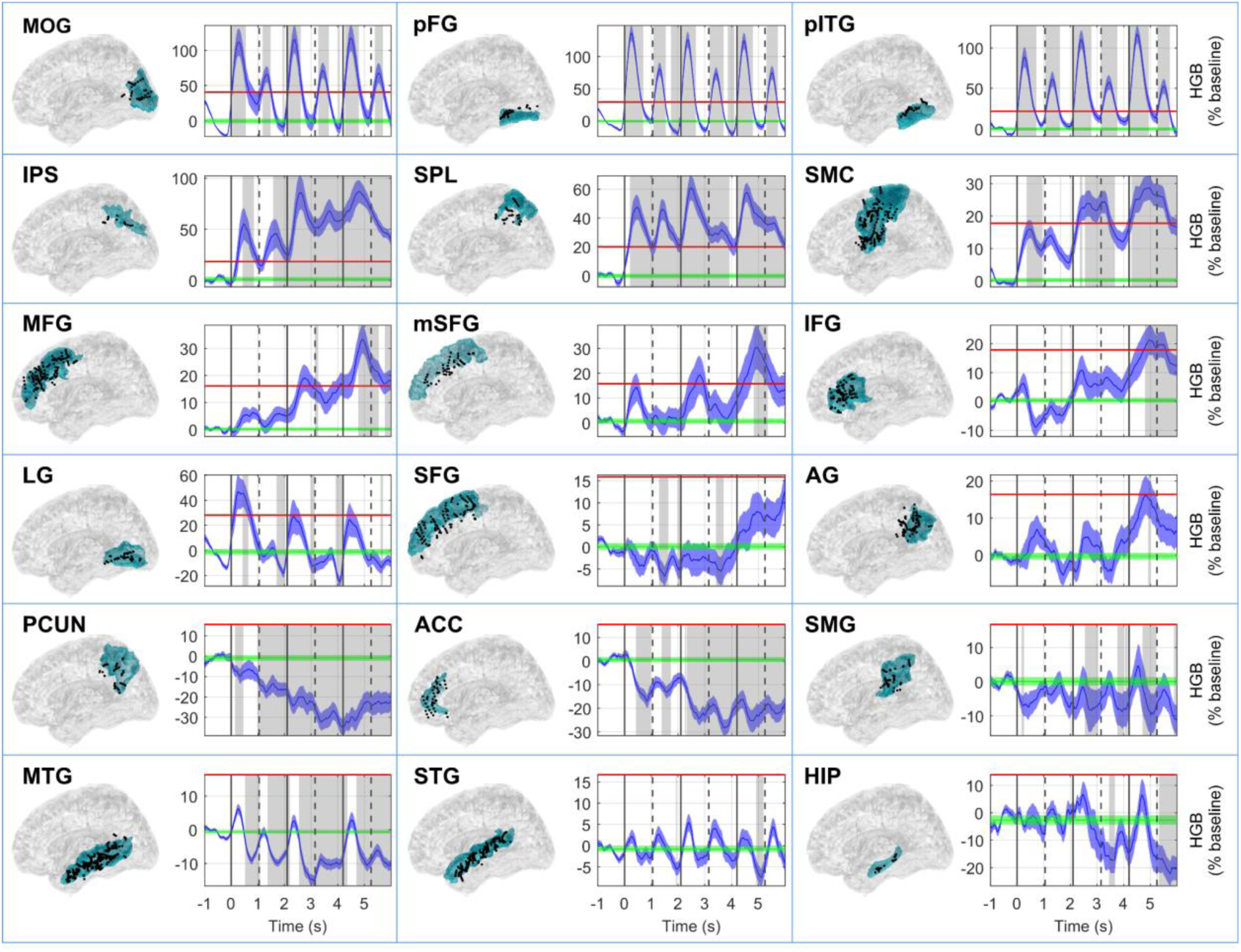
HGB activity in ROIs of the arithmetic network. Each box representing one ROI is composed of two subplots: a topology of the channels and the HGB power (52–120 Hz). The name of the ROI is indicated by the title in the upper left corner of each box. In the topology plots, the position of the channels (i.e., the centre of the bipolar pairs) is shown as a projection on a transparent brain template (Colin27), from left lateral view. Note that we projected channels from both hemispheres. Due to the normalization, projections and the channel inclusion criteria, which required that at least one of the bipolar contacts be localized in the ROI, some channels may appear slightly outside of the ROI. The selected ROI is highlighted in blue. The time-resolved (x-axis) HGB power was computed as the grand-average across all trial-averaged HGB activities of iEEG channels assigned to that ROI, which were baseline-normalized (z-scored) and expressed in percentages of the baseline distribution (y-axis). Vertical lines highlight the times of stimuli presentation (solid lines for operands and dashed lines for operators) in the sequential calculation task (first operator at time *t* = 0 s). Baseline HGB distribution, taken in the time interval from -1.00 s to -0.25 s, is depicted in green (mean ± 5*SEM). The threshold for selection of the highly activated areas (set as 2𝜎 of the baseline distribution) is indicated by a horizontal red line. Significant changes in HGB power from baseline are highlighted in grey (P < 0.001, Wilcoxon rank-sum test, false discovery rate (FDR) corrected).

Direct comparison of mean HGB activity (computed during operand or operator periods) among N1, N2, N3 and between operands O1, O2 (Fig. 5) revealed multiple statistically significant differences (P < 0.05, two-sided sign test, FDR corrected). We would like to highlight several notable observations. There were statistically significant increases (1) in the parietal areas, the IPS (but not in SPL) or AG; (2) in the frontal regions (SMC, MFG, IFG). There were also significant decreases: (1) In the extended DMN (PCUN, MTG and HIP), a network known to be deactivated during cognitive tasks requiring increased external attention demands. (2) In the VOTC (MOG, pFG), which is more difficult to interpret, but could be related to higher visual saliency of the first stimulus at the beginning of the equation.

**Figure 5:**
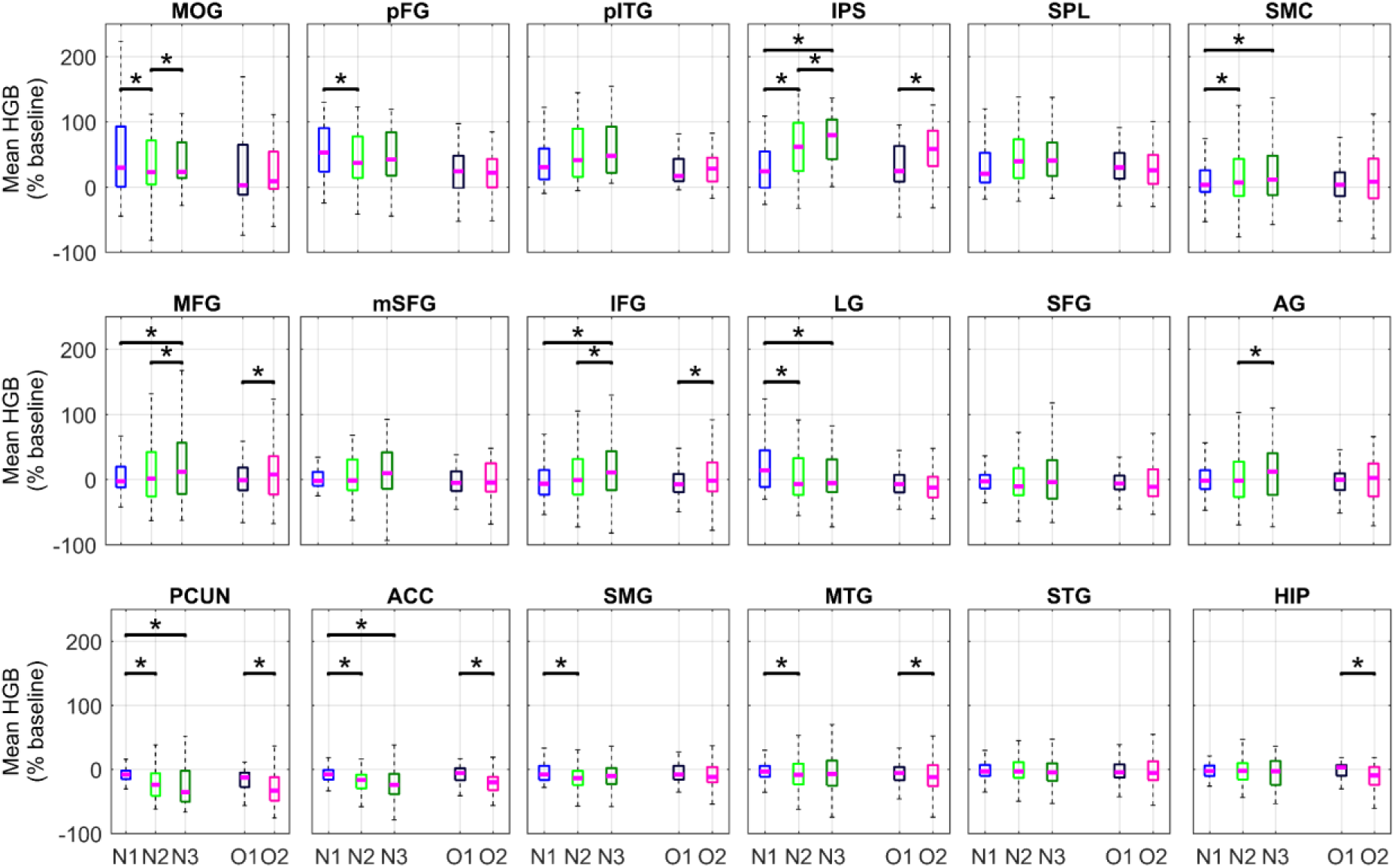
Comparison of mean HGB activity during the operand and operator periods. Each subplot represents one ROI of the arithmetic network indicated by the title (same conventions as in Fig. 1). For all channels assigned to each ROI, we computed the mean HGB responses across the operand (N1–N3) and operator (O1, O2) periods. We compared the relative changes (increases or decreases) of the channels’ mean HGB activities using the paired, two-tailed sign test. In each ROI, we tested: N1 vs. N2; N1 vs. N3; N2 vs. N3; O1 vs. O2. A star indicates significance of difference (P < 0.05, FDR-corrected for multiple testing).

### 2.3 Temporal order of HGB activations in the arithmetic network

To gain insight into the cascade of HGB activations–which may be informative about the information flow and processing within the arithmetic network–we selected the highly active regions and ordered them based on the latencies of their maxima for each operand presentation period (Fig. 6). The highly active regions in each period were defined as those crossing a set threshold (set as 2𝜎 of the baseline distribution; see horizontal red lines in Fig. 4). The HGB activity investigated in all three operand periods (N1–N3) were identical to those in Fig. 4, here only cropped to the shorter time intervals. In each operand period, we defined the local maximum of the HGB activity.

**Figure 6:**
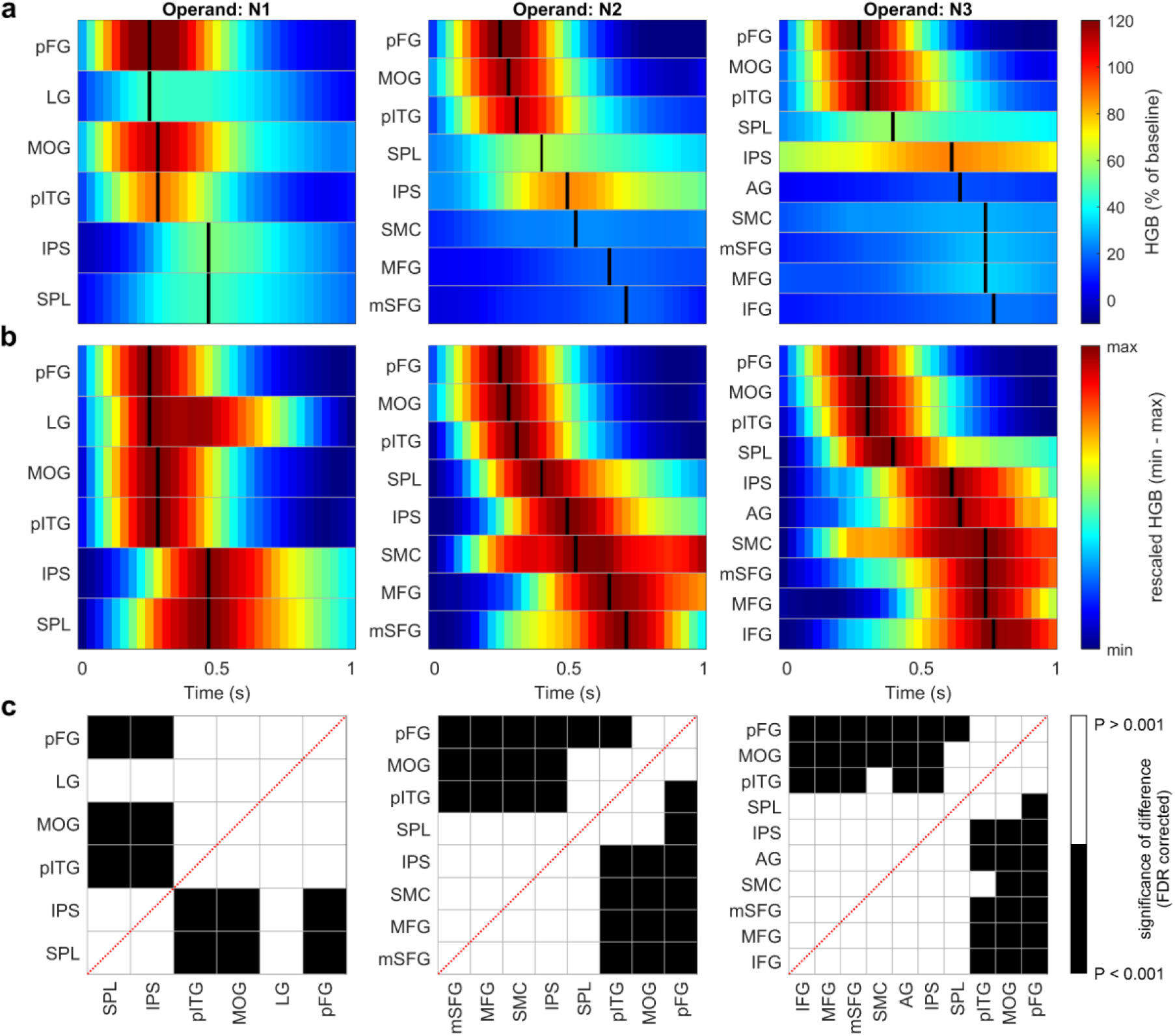
Temporal order of the HGB peaks of the arithmetic network regions. We selected those regions of the arithmetic network with highly pronounced increases from their baseline and sorted them by the latencies of their maximal values in each operand period (columns). **(a)** The HGB activity was computed against the baseline (same as in Fig. 4, but here colour-coded). Each row in the subplot represents one ROI of the arithmetic network (indicated by the title on the y-axis). The time (x-axis) is relative to the appearance of the operator (t = 0 s). Short black lines in each row mark the maximum HGB activation. **(b)** Same as in a, but with HGB values rescaled to the minimum and maximum of the ROI activity in each operator period, to more clearly illustrate the relative HGB increases of each ROI in each period. **(c)** Each chessboard represents the statistical comparison for all ROIs, showing the significance of differences in distributions of maxima latencies obtained from the single-channel, trial-averaged HGB (black: P < 0.001, Wilcoxon rank-sum test, FDR corrected; else white). We observed a reproducible pattern of cortical activation cascade underlying arithmetic network processing, with the VOTC activated first, followed by parietal and frontal areas, with the latter becoming more active during the computation-related periods (N2, N3).

There was a notable difference between the HGB responses after the presentation of the first operand (N1)—related only to the recognition of numerical symbols—and the later operands (N2, N3), which are also related to calculation processes alongside other domain-general cognitive processes. For N1, we observed the earliest peaks in the MOG and VOTC (LG, pITG, and pFG), followed by the lateral parietal areas (SPL and IPS), with significant differences in their latency maxima distributions (P < 0.05, FDR corrected). For N2 and N3, there were similar patterns to N1, with additional, later peaks in the SMC and frontal areas (MFG and mSFG), possibly related to increased attentional and working memory load during the sequential stimulus presentation.

### 2.4 HGB activity for different numerical formats, operators and problem difficulties

We also investigated the HGB activity for the different numerical formats in the ROIs of the arithmetic network, addressing the extent to which the responses were abstract or format-dependent (Suppl. Fig. S4). We found an overall high similarity among the different formats (CC = 0.71 ± 0.05, mean correlation coefficient (CC) across all formats and ROIs), especially in the highly activated areas (CC > 0.80), justifying the analyses above that disregarded numerical format information. There were, however, several notable observations: (1) HGB responses in the VOTC were significantly higher for fingers than for the other formats, which could be ascribed to the higher visual complexity of the former. (2) The pITG, supposed to contain the number form area, was not found to be selective for Arabic digits. (3) There were no significant differences and a high correlation in HGB activity (CC = 0.93) among the numerical formats in the IPS, a key area for the processing of numbers and their semantic representations, suggesting that it uses an abstract, format-independent encoding of arithmetic operations.

To delineate the potentially different strategies subjects could employ during calculation (fact retrieval during addition and procedural strategies during subtraction), we compared the mean HGB activity following the different operators (i.e., plus and minus) in the easy task (results < 10). Our results did not reveal any statistically significant differences in the mean HGB activity computed across the periods of the operands (N2 or N3) following the different operators (Suppl. Fig. S6). These results could indicate that a similar strategy was used for both operations (see Discussion for more details).

To tackle the issue of differential HGB activity as a function of problem difficulty, we computed HGB responses following those operands during which the calculation did (or did not) cross the decade in the “Arabic difficult” trials. In these trials, the decade crossing occurred once in each trial with equal probability at operands N2 and N3 (e.g., 45+6+3 crosses the decade, 50, at operand N2, while 45+3+6 crosses it at N3). We found only one region, the IPS, where the HGB activity was significantly higher for calculations that crossed the decade as compared to when they did not (see Suppl. Fig. S7), further highlighting the key role of the IPS in arithmetic processing.

### 2.5 Functional connectivity of the arithmetic network: phase-locking value

Next, we investigated FC dynamics within the arithmetic network. We applied a graph-theoretical approach, where the different arithmetic network regions were considered nodes and the time-resolved strength of the interactions represented the edges. We used the phase-locking value (PLV)^45^ as the measure of the FC based on the assumption that the phase of slower-frequency oscillations (e.g., delta: 0–3 Hz; theta: 3–7 Hz) modulates the activity of the high-frequency bands (e.g., HGB)^46^. The modulation has widespread implications in different fields of neuroscience^47^, including the connectivity studies based on iEEG^48–51^. Furthermore, the PLV is not explicitly dependent on iEEG signal amplitude, but only on the signal phase; this analysis is therefore qualitatively different from the above activation analysis, which was based exclusively on the iEEG signal power regardless of its phase. The time-resolved strength of the PLV between two different regions of the network was computed as the mean across all respective channel pairs on a single-subjects level (i.e., without mixing channels from different subjects; see Suppl. Table T2), against the baseline distribution. We used all trials with correct answers, regardless of the numerical format or operator sign—a simplification, which can be partially justified by the relatively large similarity of the HGB responses for the different numerical formats (Suppl. Fig. S4) or operators (Suppl. Fig. S6) and also by the need of a large number of trials to estimate the PLV reliably.

The PLVs were most pronounced in the lowest frequencies, the delta (0.1–3 Hz) and theta (3–7 Hz) bands (see Suppl. Fig. S8). The “global” network connectivity (i.e., average over all channel pairs for all combinations of regions) was significantly increased with respect to the baseline values (Fig. 7a) and revealed three distinct peaks after the presentation of the operands, while the PLV peaks for the operators were much less pronounced, especially in the delta band. Of note, the theta PLV preceded the delta PLV. More specifically, the theta PLV “offset” times (defined as the minimal slope of the theta PLV during the operand periods; downward-pointing triangles in Fig. 7a) preceded the times of delta PLV peaks. This temporal difference points towards their differential roles in network synchronization and activation.

**Figure 7:**
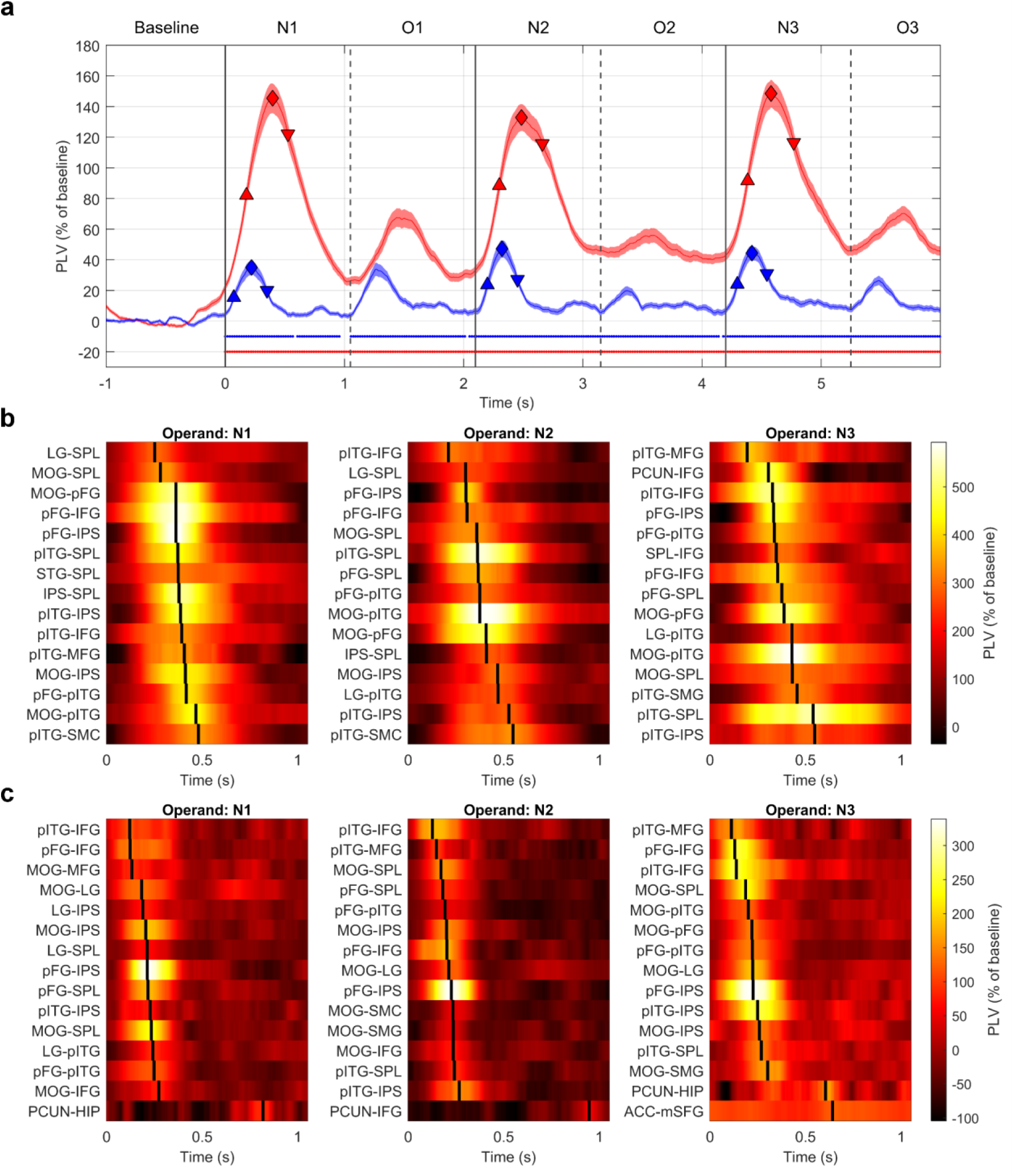
Temporal profiles of functional connectivity within the arithmetic network. **(a)** Mean PLV (mean ± SEM across all channel-pairs of all ROIs) for two different frequency bands: delta (0.1–3 Hz; red color) and theta (4–7 Hz; blue). For each operand (N1–N3), we defined three time points: onset (defined as maximal slope; upward-pointing triangle), peak (diamond) and offset (defined as minimal slope; downward-pointing triangle). Dots at the bottom of corresponding color indicate significance of difference against the baseline values distribution (P < 0.05, Wilcoxon rank-sum test, FDR corrected for multiple testing over different time points). Vertical solid or dashed lines represent the time of the presentation of the stimuli (operands N1–N3 or operators). **(b)** In each operand period (N1–N3; figure columns), we selected 15 pairs of arithmetic network ROIs based on their maximum PLV value in the delta band, and ordered them according to the latencies of the maxima. Temporal profiles (x-axis) of PLVs between the 15 strongest ROI pairs (rows of the subplots; names indicated on the y-axis). **(c)** Same as in b, but for the theta band.

We also investigated the temporal sequence of the FC’ strength changes by ordering the latency of the PLV maxima (Fig. 7b, c; Suppl. Fig. S10) and onset (Suppl. Fig. S9 for delta band and theta band). We selected 15 ROI pairs with the strongest connections (separately for each operand-related period). Similar to the HGB analysis above, the maximal PLV value had to exceed a threshold set as 3𝜎 of the baseline distribution. This ensured that we investigated only highly pronounced increases in the FC rather than noisy fluctuations in the PLVs.

### 2.6 Patterns of functional connectivity within the arithmetic network

As a final synthesis of the previous results, we illustrate the HGB activations and PLV connectivity strengths separately in the theta and delta frequency bands in time-resolved diagrams (Fig. 8; see also a full animation in supplementary videos V1–V4). PLV connectivity showed a broadly consistent pattern. Some differences were noted between operands N1 (mere digit recognition) and N2/N3 (active calculation), as well as between the theta and delta frequency bands. Generally speaking, connectivity patterns were more extensive during the active calculation stage, especially in the theta band. To illustrate these connectivity patterns, we chose three time points for each operand based on global PLV connectivity (Fig. 7a): the onset, peak, and offset (see Methods for more details).

**Figure 8:**
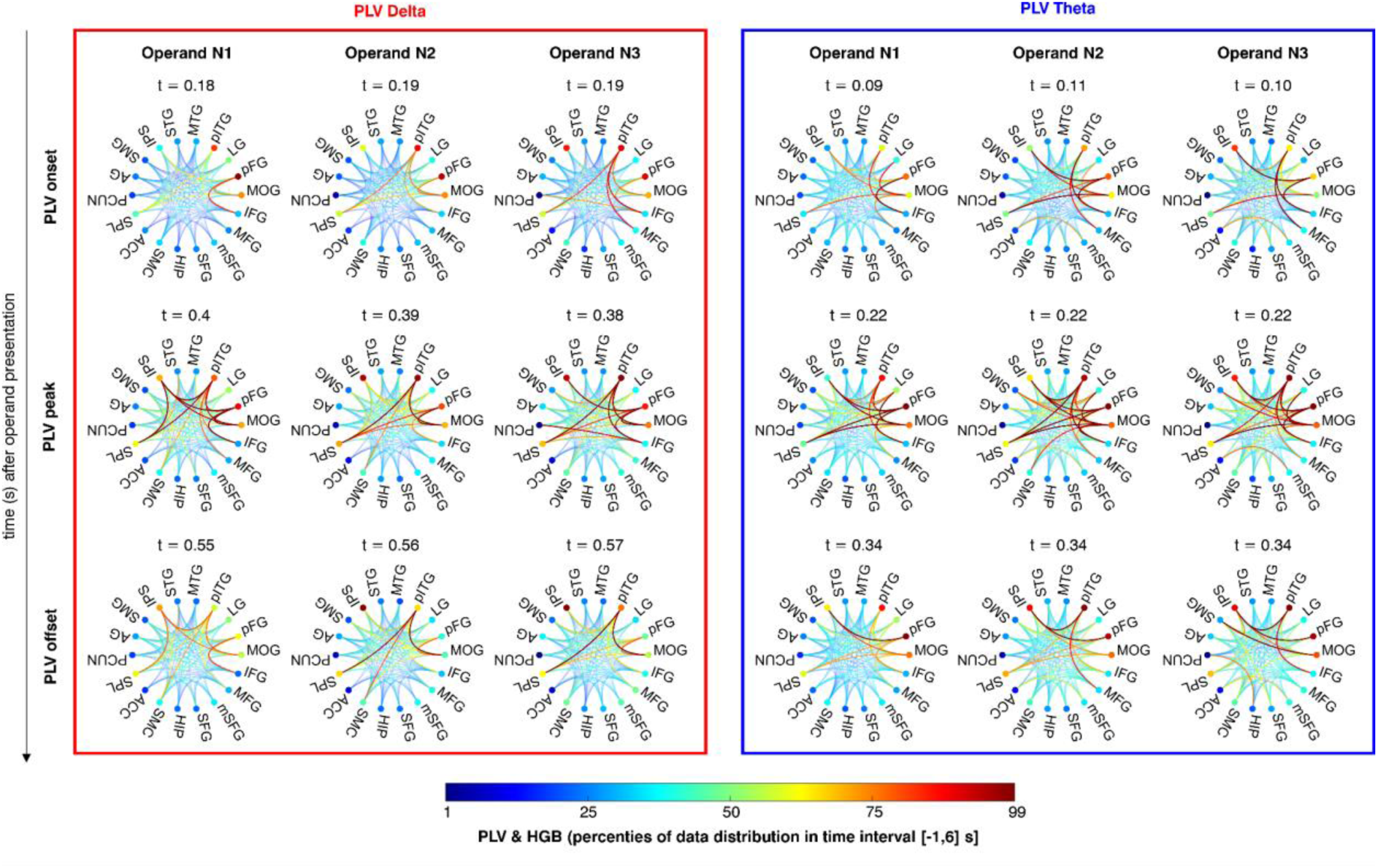
Patterns of HGB activation and PLV connectivity of the arithmetic network. Each scheme represents a unique state of HGB activation and connectivity of the arithmetic network during the sequential calculation task. The dots on the circumference of the scheme depict ROIs with normalised HGB activation (colour-coded, where minimum/maximum was set to the 1–99 percentile of all HGB values across all ROIs for time 0–6 s), and the lines connecting ROIs represent normalized PLV connectivity strength (colour-coded, with minimum/maximum set to the 1–99 percentile of all PLV values for time 0–6 s). Columns represent the time periods of the operands N1–N3. Rows represent either PLV onset, peak or offset (see Fig. 7a), with times indicated above each scheme. Rows represent the selected time points (indicated on the left). A pronounced and reproducible pattern of activation and connectivity emerged around 200 ms after the presentation of the operands for theta band and 400 ms for the delta band, with notable differences among the operands.

At PLV onset (the maximal slope), we observed early connectivity between VOTC-frontal regions and VOTC-parietal regions in both frequency bands (apart from operand N1 in the delta band). Interestingly, the frontal connectivity changes occurred prior to respective increases in HGB activity. As the PLV reached its peak, connectivity patterns became more extensive, including all VOTC, parietal (IPS, SPL), and frontal areas (IFG, MFG). Both bands showed additional VOTC-SMC and VOTC-SMG connections. At N3, we also observed IFG-PCUN connections in the delta band, and ACC-mSFG connections in the theta band. During PLV offset (the minimal slope), connectivity patterns started to diminish. In the delta band, connections centered on the pITG, which was linked to the MOG throughout, to the IPS at N1, and to the SMC and SPL during active computation, especially at N2. The theta band showed a strong pFG-IPS connection at N1, with additional pITG-IPS, pITG-IFG, MOG-SMG, and PCUN-HIP connections developing only later during the computation stage.

The aforementioned connections diminished after 800 ms in the delta band (apart from pITG–SPL during N3 lasting until 900ms), and after 500 ms in the theta band (see Supplementary videos V1–V4). Interestingly, the PLV connectivity in the theta band showed a less pronounced, secondary peak at around 700 ms after the presentation of each operand (see Fig. 7a - smaller second peak of PLV during operands N1-N3, and Fig. 7c); this included connections involving HGB deactivated regions, particularly frontal regions (ACC-mSFG, IFG-pITG) and the HIP (HIP-SMG, HIP-pFG, HIP-PCUN).

## Discussion

This study provides insights into the temporal dynamics of activity and connectivity patterns within the putative arithmetic network during a sequential arithmetic task. Summarizing the main results, we observed that: (1) The VOTC areas (pITG and pFG) manifested short, transient activations (Figs. 3 and 4), serving as a key hub in the network, presumably relaying the visually processed information to the other network ROIs. Contrary to our expectations however, the pITG did not show signs of specificity for Arabic digits, with its responses being instead largely format-independent (Fig. S4). (2) The lateral parietal regions (IPS and SPL) showed a sustained, gradually increasing activity over time (Figs. 4 and S5), indicating their more direct involvement in computation itself. Additionally, the IPS reflected problem difficulty effects (Fig. S7) with no differences between numerical formats (Fig. S4), supporting its role in the manipulation of abstract magnitudes. (3) The frontal areas (IFG, MFG, and mSFG) were activated during more demanding periods of the task (Fig. 4), possibly related to the allocation of resources for attention or working memory. (4) Areas of the default mode network (PCUN, MTG, and HIP) gradually deactivated (Figs. 4 and S5), presumably to ensure proper allocation of the brain’s computational resources. (5) The temporal order of HGB activity started in the MOG and VOTC, before activation of lateral parietal and—during calculation periods—sensorimotor regions, and finally frontal areas (Fig. 6). (6) The VOTC showed no signs of specificity for Arabic digits, with signs of general format independence noted in the IPS, supporting the idea that this region hosts abstract representations of magnitude (Fig. S4). (7) The VOTC regions (pITG and pFG) elicited early (100–200 ms after the presentation of each operand) FC with frontal regions (IFG and MFG; Fig 7b, c and Suppl. Fig. S9). (8) The theta PLV peaked earlier than the delta PLV (Fig. 7a). (9) The FC pattern during periods of calculation (N2/N3) were more extensive than during the mere number recognition period (N1) (Fig. 8). (10) A stable and reproducible FC pattern emerged among the major hubs of the arithmetic network (200–400 ms after the presentation of each operand) with interpretable differences between the operands (Fig. 8).

We stress that the results of this study pertain to a comparison of a sequential arithmetic task with an inter-trial baseline, not a control cognitive task. As such, our findings cannot be interpreted as arithmetic-specific. While considering cognitive strategies, problem difficulty, and other within-task contrasts may help approximate the type of activity captured, the absence of a control task ultimately precludes us from distinguishing neural activity specific to arithmetic from that related to domain-general functions, such as working memory, attention allocation, visual processing, or reading and verbalization^52^. In the arithmetic literature, reference conditions like the resting state^53–55^ or fixation period^36,46,56^ are instead used to capture general task-related activity, isolating it only from ongoing, task-irrelevant background processes. While studies with control tasks typically show at least partial specificity within the arithmetic network^8,37^, the issue remains an important topic for further research. Acknowledging this limitation, we discuss our results below based on their anatomical location (from occipital, temporal, parietal, to frontal), alongside several other interesting observations, study limitations, and open research questions.

The VOTC showed the earliest activations, corroborating previous evidence^3^. The pITG is often placed among key numerical cognition structures based on research suggesting its specificity for numerical symbols or Arabic digits^4,5^. Our results (Fig. S4) align more closely with counter-arguments indicating a lack of format dependence within the pITG^7,57^. Similarly to other VOTC regions, the pITG showed a preferential response only to fingers, likely due to the higher visual complexity of these stimuli. Notably, our results also indicated that the VOTC’s role in numerical processing may go beyond the mere visual processing of numbers. In particular, our finding of strong PLV connectivity with both the VOTC and parietal regions (Figs. 7, 8) indicates that the occipital areas may be involved in the continuous updating of the visually presented numerical information, in cooperation with the main numerical hubs. We also analysed data from the LG and found that the HGB activity profile was almost entirely specific for operands, but not operators, and that its peaks significantly decreased across the operands (Fig. 5). These novel observations may support a role in symbolic number processing, as reported in several fMRI studies^8,26,27,58^.

Our results regarding the VOTC support its hypothesized role as a visual number code hub^59^. We observed a pronounced peak activity shortly (200 ms) after the presentation of each operand. Notably, strong and highly correlated HGB responses were detected in both the pITG and pFG (Fig. 4), further fueling the ongoing debate concerning the localization and specialization of the putative number form area^2,9,26^. However, there were some notable differences. The pFG peaked earlier than the pITG (although the differences in the maxima latency distributions were significant only after the N2 operand). Both pITG and pFG demonstrated early and robust PLV connectivity with the parietal regions (SPL, IPS), in line with a recent report of the VOTC influencing the lateral parietal cortex^21^. An interesting and rather unexpected observation was the early onset of high PLVs between VOTC and frontal areas (Figs. 7, 8, and Suppl. Fig. S9).

In contrast to the VOTC, the MTG and STG were much less activated, even though several fMRI studies have previously shown their activations during arithmetic^5,10,19,26,27^. Their transient activations after the presentation of each operand were followed by deactivations (Fig. 4). Likewise, the HIP showed only mild engagement based on its HGB activity, but demonstrated an interesting late PLV connectivity in the theta band with SMG, pFG or PCUN^21^.

Our findings in lateral parietal regions (IPS and SPL) confirm their prominent role within the arithmetic network and partially align with previous reports. In particular, the IPS exhibited the most prominent HGB activity, characterized by the highest amplitude, longest duration, and most consistent and significant increase across operands (Figs. 4, 5 and S5). High correlations in HGB activity were noted between numerical formats with no significant differences between them, suggesting that the IPS is largely format-independent. Overall, these findings support the region’s central role in the analogue representation of numerical magnitude and its reliable engagement across various numerical tasks, including arithmetic^10–12,53^. Compared to the IPS, the SPL was activated slightly earlier for all operands (although the differences in peak latency distributions were not significant), and the magnitude of change from the baseline was less pronounced, with no significant difference among operands N1–N3. This suggests a supportive role during arithmetic processing, possibly related to topographical representation of quantities^54^ or attention allocation^55^. The activity of regions in the inferior parietal lobule (AG and SMG), associated with the verbal numerical code ^8,11^, was less pronounced compared to the robust activity increases of the IPS or SPL and was consequently, more difficult to interpret (Fig. 4). These regions are part of the association parietal cortex and thus contain heterogeneous subpopulations with distinct functional roles, which are likely obscured in grand-averaged measure of activity.

Interestingly, we observed robust HGB deactivation in the PCUN, MTG, and HIP (Fig. 4). These findings stand in contrast to evidence of their involvement in arithmetic and symbolic number processing in neuroimaging studies^5,10^. However, they do support a role for these structures within the default mode network, which is typically suppressed during externally directed cognitive tasks such as maths^20,43,56^ or attention allocation^49,60^. Mental effort can also perturb activity in a partially overlapping brain network involved in autonomic control^61,62^. This may explain the additional HGB deactivation observed in the ACC, contrasting previous fMRI findings ^9,20,27^. Notably, and as also seen here for the ACC, such deactivations can co-occur with increased theta activity (Fig. 3), suggesting that they reflect fluctuations in working memory demands rather than a simple shutdown of a default process^61,63^.

Our results in the frontal lobe revealed that HGB activity showed a gradual, significant increase through the task in the SMC, MFG, and IFG (Figs. 4, S5), supporting the involvement of these regions in calculation and/or associated cognitive functions like working memory and decision making^1,10^. The activity in SMC is consistent with prior studies highlighting the importance of finger-counting strategies during the development of numerical cognition in children, suggesting that these circuits also remain engaged during arithmetic processing in adults^10,12,34^. Among the frontal regions, the mSFG and MFG showed the overall highest HGB activity, followed by the IFG. The prominent simultaneous engagement of these regions in particular may reflect the increased cognitive demands of our sequential task and the associated attention and working memory load. These findings align with the updated triple code model proposed by Arsalidou and Taylor^2^, suggesting that frontal areas contribute differentially depending on the task difficulty.

The robust frontal activations, combined with the weaker response of some temporal regions (MTG, STG) and the inferior parietal lobule (SMG, AG), suggest that the present study captured primarily procedural, as opposed to fact-retrieval strategies. We speculate that this may be due to the sequential presentation of stimuli, which may encourage fact-retrieval strategies less than simultaneous designs. For example, Pinheiro-Chagas et al. observed a higher rate of deactivations across the inferior parietal lobule during sequential compared to simultaneous tasks^43^. Task-related factors may also explain the muted response of medial temporal structures like the HIP, found crucial for mental arithmetic in a previous iEEG study using a simultaneous task^21^. In this respect, the fact that we also observed no significant HGB activity differences between addition and subtraction may seem less surprising. We note, however, that while many arithmetic models predict that the repeated exposure to single-digit addition results in a progressive shift from procedural strategies to fact retrieval^15^, they have not gone uncontested. More recent research suggests that simple addition may instead rely on time-efficient counting strategies^17,18^, providing an additional explanation for our findings.

Notably, while frontal regions showed increased activity during calculation periods, they did not respond to problem difficulty, operationalized here as the cross-decade factor. The IPS alone showed a heightened response, supporting its role as a “core quantity system”^11^. Nonetheless, the specificity of the IPS to calculation can be debated; for example, previous iEEG studies documented problem difficulty effects not just in the IPS, but also the SPL and other arithmetic network nodes^3,28^.

Several limitations, inherent to the iEEG method^68^, stem from the uneven and sparse distribution of the intracranial electrode contacts in the brain and a relatively small and heterogeneous cohort, consisting of epilepsy patients. These limitations impose certain challenges for iEEG data processing, statistical testing for significance of difference between two ROIs with different sample sizes, and especially for interpreting the results. In order to arrive at a holistic description of arithmetic network dynamics, we averaged the results obtained from single-channel activity (or channel-pair connectivity) in each ROI^69^, rendering the activity of each ROI homogeneous. Ignoring the differences within each ROI in this way is likely an oversimplification, as some studies have reported a degree of heterogeneous responses and interactions^3,6^. We also did not take into account lateralization, regarding heterogenous handedness and implantation schemes of the subjects. This might have resulted in bias, particularly in some regions with predominantly left-lateralized activity in numerical cognition tasks, such as the AG^19^.

FC results, assessed here through PLV, should be interpreted with great caution^70^. While some of the typical pitfalls, such as common reference, volume conduction, or sample-size bias problems, were mitigated by use of a bipolar reference or comparison against baseline values distribution (using the same number of trials), other issues should be noted. Namely, the common input problem, which is especially hard to tackle if one does not observe all nodes simultaneously, may result in spurious positive PLV results between ROI pairs. Furthermore, one should refrain from interpreting the connections as being necessarily direct, as they can be mediated by other, unobserved nodes of the network. In our PLV analysis, we concentrated only on phases of a low-frequency bands: delta (0.1–3 Hz) and theta (3–7 Hz). While we observed the most pronounced differences (compared to baseline values) in these bands, the optimal frequency range could be specific to each ROI pair, as recently reported^50^. Nevertheless, low frequencies have been implicated in facilitating long-range communication in the brain^49,71^.

The PLV analysis most likely indicates the early, initial synchronization of the network, the probable mechanism is theta-gamma coupling, in which the phase of the low-frequency (here 0.1–7 Hz) oscillations drives the high-frequency (> 50 Hz) amplitude^72^, in turn facilitating task-related spiking activity^73^. In this light, it is of note that the theta PLV offset systematically preceded the delta PLV peaks in all three operand periods (N1–N3), implying their differential roles in genesis of cortical activation. The lack of PLV connectivity, particularly in the delta band, at the later stage (> 800 ms after the presentation of the operands) could be considered surprising, given the high HGB activity in some ROIs (IPS, SPL, MFG, mSFG, and IFG). The reported PLVs could thus be related to higher amplitudes of the iEEG signals in the delta and theta bands after the presentation of the operands (event-related potentials), which in turn allow for a better phase estimation in the otherwise noisy signal^74^. Due to the sparse spatial sampling of the iEEG electrodes, some ROI pairs were not sampled at all, and connectivity between them could not be reported. This included pairings between the ACC, SFG, mSFG, and the occipital/temporal/parietal areas (see Suppl. Table 2). We also note that we applied only a limited set of analyses regarding FC. Future studies could build on this work by investigating directed (or effective) connectivity measures or disrupting network dynamics through transcranial magnetic stimulation, for example.

In conclusion, our study confirms and extends previous findings from iEEG data on a putative arithmetic network^3,6,34,35^, reports that remain relatively scarce given the unique and invasive nature of this method. In particular, we confirm the robust activity of the major hubs of the arithmetic network, the VOTC (pITG, pFG) and lateral parietal areas (IPS, SPL), but also note activity in the frontal regions that increases with the task demands. We also offer new insights about the FC within the arithmetic network, highlighting early FC patterns between VOTC and frontal areas as well as robust and reproducible patterns emerging 100–500 ms after the presentation of each operand during a sequential arithmetic task. Altogether, these results can be used to guide and refine future arithmetic network models or electromagnetic interventions.

## Methods

### Participants

Twenty subjects (eight females, average age 36.2 years, range 18–63) with pharmacoresistant epilepsy were included in the present study. During pre-surgical evaluation, they underwent iEEG video monitoring and were implanted with stereo EEG (SEEG) electrodes to assess the epileptogenic zone. The duration of SEEG implantation typically varied from 7–14 days. The implantation scheme depended solely on the clinical hypothesis. Subjects participated voluntarily in the study after signing informed consent. The study was approved by the ethical committee of Motol University Hospital (EK-1340.27/20), Prague, Czechia. All ethical regulations relevant to human research participants were performed in accordance with the Declaration of Helsinki. Electrode implantation and subject details are summarized in Suppl. Table T1 and Suppl. Fig. S1.

### Sequential arithmetic equations task

The task consisted of multiple trials (repetitions) of different, sequentially presented, three-operand, arithmetic equations (Fig. 2), for example: 6 + 3 - 2 = 8 (yes/no). Each trial began with a fixation period, followed by the sequential appearance of operand N1 (number), operator O1 (either plus or minus), operand N2, operator O2 (either plus or minus), operand N3, equals symbol, and result. The result was correct in 50% of the trials, and in the remaining trials differed by one (either smaller or larger). The participants’ goal was to determine whether the result was true or false and press a corresponding button on the gamepad (Logitech F310): green or red button for a true or false answer, respectively. Furthermore, the subjects could also press a blue button to indicate that they did not know the answer, to avoid guessing. For this reason (and also to retain most of the unique iEEG data), we did not exclude subjects whose overall performance approached the 50% chance level. Here, we will refer to the presented operands, operators and the fixation cross as “stimuli”.

All stimuli were presented for 1.05 s, with the exception of the fixation period, which was randomly varied between 1.05 and 1.55 s. The subjects had 5 s to provide their answer before the trial was timed-out. Based on the answer (or a time-out), feedback was provided: correct answers triggering a smiling cartoon face together with a “hooray” sound; incorrect, “don’t know” answers, and time-outs triggered a sad face and a “boo” sound. The following trial started (with the fixation cross) after the feedback. The task was implemented in PsychToolBox 3.0^75^ and the presentation of each stimulus was synchronized with the iEEG recordings via a parallel port-connected trigger channel to enable trial extraction.

There were two difficulty levels: “easy equations”, with answers that did not exceed 10, and “difficult equations”, with answers greater than 10, for example: 54 - 7 + 2 = 48 (yes/no). In the “easy equations”, the operand stimuli were selected from the following “number categories”: Arabic numbers, dice symbols, fingers, and words (e.g., “six”). Every equation presented only one stimulus type (fingers and dice symbols were not mixed, for example). In the “difficult equations”, the operands were presented only as Arabic numbers. The proposed result was always presented as an Arabic number. Other stimulus types and tasks were also conducted, but were not analyzed in the current study, these included auditorily presented numbers and letters presented both visually and auditorily. All stimuli were white and presented on a dark grey background. Subjects, situated at the neurology ward, were sitting in their beds while a PC screen was placed in front of them at their chest level, approximately 1.5 metres from their eyes.

The task was first explained to the subjects, and they were then able to practice until they were clear about the task. The task was split into multiple recording sessions. A session could be either the “easy equations” (10 repetitions for each stimulus type) lasting around 6 minutes, or the “difficult equations” lasting around 2 minutes. We purposefully split the easy and difficult conditions, so each subject would have the option to stop this more challenging part at any point at which they did not feel comfortable. We ran 4–5 sessions (depending on the subject’s preference), gathering at least 40 trials (equations) for each stimulus category. There were pauses in between the sessions, the length of which depended on the subjects. Subjects also had the option to divide the task into multiple (typically two) days.

### iEEG data recording and preprocessing

iEEG data were recorded by SEEG electrodes (DIXI Medical) sampled at 2048 Hz by Natus Quantum amplifiers, with the reference and the ground electrodes placed at the depth of the white matter. SEEG contacts located in the seizure onset or irritative zones—delineated by experienced neurologists—were rejected from further analyses to avoid including potentially pathological brain areas. The iEEG signal was first low-pass filtered (8th order Chebyshev Type I infinite impulse response filter, cutoff at 400 Hz, zero-phase shift) to prevent aliasing and downsampled to 512 Hz to reduce the computational memory load. The iEEG signal from the remaining contacts was visually examined to reject broken channels with artefacts. We then applied bipolar referencing of the signal, between every contact and its closest neighbour at each SEEG electrode shank, starting from the deepest contact on the electrode. The aim was to increase the local specificity of the iEEG signal, eliminate the influence of the common reference, and attenuate the influence of more distal neural sources (as recommended in^76^ and as we have done in our previous studies^77–79)^. We refer to the signal measured by bipolar-referenced SEEG contacts simply as a “channel”. All channels were high-pass filtered to remove slow drifts (0.1 Hz cutoff, Butterworth, 6th order, zero phase shift) and notch-filtered at 50 Hz and its harmonics (100, 150 Hz) to reduce line noise (e.g., 48–52 Hz stop band, Butterworth filter, 6th order, zero phase shift). We also marked the inter-ictal discharges in the iEEG data using the epileptic spike detector^80^. The detected epileptic spikes (and the 0.5-s time window centered on the spikes) were rejected. These preprocessed iEEG data were subject to further analysis.

### Channel Localization

Localization of SEEG electrode contacts was assessed by co-registering post-implantation CT scans and pre-implantation MRI; positions were also later verified on a post-implantation MRI, to exclude potential shifts of brain parenchyma due to oedema or bleeding. Every SEEG electrode contact was automatically detected^81^ and visually assigned to anatomical regions based on anatomical gyrification specific to each subject by an experienced neurologist, according to an anatomical atlas^82^. Later, the MRI scans were normalized to MNI space using SPM12 (statistical parameter mapping^83^) to obtain MNI coordinates of the SEEG contacts.

### Regions of interest (ROIs) of the putative arithmetic network

From the anatomical localization of each SEEG electrode contact, we assigned the (bipolar) channels to ROIs, which were selected based on previous neuroimaging, electrophysiological, and neuropsychological reports. For the assignment, we demanded that at least one of the two bipolar contacts be located in the ROI, to preserve the maximum number of channels. We also included channels located on the boundaries of grey and white matter, because channels even a few centimetres away from the source have been shown to record high fidelity signal^84^.

We included the following 18 regions with a reported role in the putative arithmetic network: the middle occipital gyrus (MOG; corresponding to lateral occipital gyrus “LOG” in several studies)^2^, posterior fusiform gyrus (pFG)^34^, posterior inferior temporal gyrus (pITG)^4^, lingual gyrus (LG)^8^, middle temporal gyrus (MTG)^5^, superior temporal gyrus (STG)^20^, hippocampus (HIP)^21^, intraparietal sulcus (IPS)^6^, superior parietal lobule (SPL)^19^, precuneus (PCUN)^9^, angular gyrus (AG)^85^, supramarginal gyrus (SMG)^86^, anterior cingulate cortex (ACC)^5^, sensorimotor cortex including precentral and postcentral gyri, sulcus centralis and central operculum (SMC)^2,87^, inferior frontal gyrus (IFG)^27^, middle frontal gyrus (MFG)^88^, superior frontal gyrus (SFG)^2^ and its medial part (mSFG)^20^. Of note, this network selection included all nine regions of the math-network proposed by the recent study of Pinheiro-Chagas et al.^3^.

In the cases of the FG and ITG, we decided to include only the posterior part of the gyri, as most of the numerically associated activations have been described there^4,5,57^. For both regions, we defined the posterior HIP as their anterior boundary, corresponding to the posterior–anterior coordinate Y = - 40 mm in the MNI space. We manually checked the normalized MNI Y-values for all the contacts, and then visually checked the position of each marginal contact with respect to the HIP in the individual brain before the normalization.

### Trial extraction

From the preprocessed iEEG data, we extracted trials based on the trigger-channel synchronization in the time interval from -2 to 9 s, where time t = 0 s corresponds to the presentation of the first operand N1 (see Fig. 2). We purposefully extracted a somewhat larger trial window to avoid edge artifacts during signal filtering, and cropped the trials to the time interval from -1 to 6 s for results visualization. Thus, for each subject *s*, the result of the trial extraction was a 3-D matrix of iEEG data: *D_s_*(*t*,*ch*,*tr*), where *t* refers to the samples of the trial time, *ch* to a channel assigned to a specific ROI, and *tr* to a trial. We will refer to this data set as “raw trials”.

### Arithmetic network dynamics from the iEEG signal

Our goal in this study was to describe a network-level activity and interactions, estimated here either by a relative frequency band power or a FC measure (see below). Here, we would like to explain the rationale common to both approaches, and how we derived the network dynamics from single trials at single channels measured in multiple subjects. In our view, the (bipolar referenced) iEEG channels—measuring the macroscopic local field potentials^67^—sample brain activity in different ROIs. By averaging across all trials, we get a robust estimate of the activity (or interactions) at the single channel (or channel-pair) level. The ensemble of the trial-averaged channels assigned to one particular ROI constituted the ROI-specific distribution. By averaging across all “trial-averaged” channels assigned to one particular ROI, we arrive at an estimate of the ROI-related activity. Similarly, by averaging across all channel pairs assigned to two different (non-overlapping) ROIs, we get an estimate of ROIs-related interactions. Another common feature in our analyses was that all ROI activations (or interactions) were expressed as a percentage change from the baseline (fixation cross) period.

### Spectrograms of the ROIs of the arithmetic network

To estimate the time-resolved PSD and the HGB activity, we applied a multitaper analysis with two, 0.5-s-long sliding Slepian tapers, with a time step of 31.25 ms across the raw trials (corresponding to 16 samples at sampling frequency 512 Hz). Multitapering yielded a 4-D matrix of raw PSD: *rawPSD_S_*(*f*,*t*,*ch*,*tr*), where *f* refers to a frequency bin (in the range of 0–120 Hz with 2-Hz resolution), *t* corresponds to the time of the sliding window center, the other indices are same as in the raw trial data. The *rawPSD* was log-transformed to dB (to partially compensate for the skew of the data distribution in each frequency band) and normalized to the baseline time period, *t* ∈ [-1, -0.25] s, to correct the 1/*f*-decay of the iEEG signal^90^, similar to our previous study^49^. In more detail, the normalization of the PSD, yielding *normPSD*, was computed as a z-score for each frequency bin *f* and each channel *ch*:

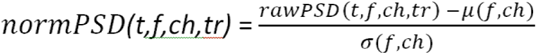

where the *μ*(*f*,*ch*) and *σ*(*f*,*ch*) are the z-score normalization factors computed from the *rawPSD* across all baseline samples of all trials. The normalized (or relative), z-scored (with respect to the baseline) *normPSD* was expressed as percentages of the baseline distribution, so that the mean baseline activity is equal to 0 %, and mean ± SD is equal to ± 100 %. Spectrograms of ROI-specific PSD were computed as a mean across all channels of all subjects in that ROI.

### HGB activity of the ROIs of the arithmetic network

The HGB activity was computed from the normalized PSD, *normPSD_S_(t,f,ch,tr)*, as a mean across frequencies *f* in the interval 52–120 Hz, excluding the notched-out harmonics at 50 and 100 Hz, yielding a 3-D matrix: *HGB_S_*(*t*,*ch*,*tr*), with same index notation as for *rawPSD*. Next, the HGB was averaged across selected trials, leading to a 2-D “trial-averaged” *HGB_S_*(*t*,*ch*), to which we will refer as a “single-channel, trial-averaged HGB”. To obtain the ROI-related HGB activity, *HGB*(*t*, *ROI*), we averaged *HGB_S_*(*t*,*ch*) across all channels *ch* of all subjects *s* assigned to the ROI. The ROI-related HGB activity was thus also expressed in percentages of the baseline distribution, because it was computed as the mean across the baseline-normalized channels. The HGB activity could be averaged across selected trials given a certain condition (e.g., numerical formats, plus/minus operators) and selected time windows (e.g., time of the operand/operator presentation).

### Indicators of HGB activity

For each channel *ch*, we computed three different indicators of the HGB activity (Fig. S5) in the 5 s time interval immediately after the presentation of the first operand (*t* = 0 s): (1) The mean of the absolute value of the HGB, *M*(*ch*) = <|*HGB*(*t*,*ch*)|>_t∈[0, 5]s_, where <> denotes the mean across time and *HGB*(*t*,*ch*) is the *single-channel, trial-averaged HGB*. (2) The duration of the significant activations (Fig. S5b), *T*(*ch*), was computed for each channel based on the single-trial values as follows: (i) The significance of difference in distribution of HGB values against baseline was computed across trials (P < 0.05, Wilcoxon rank-sum test, FDR corrected for multiple testing across all time points and channels). (ii) The duration was then computed as the sum of the significant time points. (3) The slope of the linear regression fit, *b*(*ch*), to the *single-channel, trial-averaged HGB*(*t*,*ch*).

### Interactions of the arithmetic network ROIs: PLV

To quantify the interactions and their temporal changes between the different ROIs of the arithmetic network, we computed the PLV^45^ for each sample across trials using the “filter-Hilbert” approach^49,91^. In more detail, we first divided the frequency spectrum of the iEEG signal into 4-Hz bins, with the exception of low frequencies, for which the binning followed the traditional division into delta (0–3 Hz), theta (3–7 Hz), and alpha (7–12 Hz) bands. Next, we filtered the raw trials (6th-order Butterworth filter, zero phase shift) in narrow frequency bands of 4-Hz width (i.e., 0.1–3–7–12–…–120 Hz). For each channel-pair *m* and *n* between any two ROIs and each narrow band, we computed the complex-valued analytic signal by Hilbert transform, allowing the estimation of the signal phase 𝜑 used in the PLV computation:

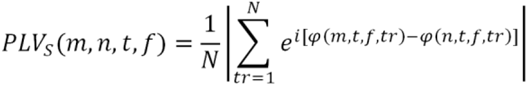

where 𝜑 is the phase of the *m*-th and *n*-th channel at time *t*, *f* denotes the narrow frequency band, *tr* is trial, *N* is the number of the trials, and *s* is the subject; this means that we computed the PLV only between channel pairs from single subject, without mixing channels from different subjects. The PLVs were normalized (z-scored) to the percentage of baseline distribution for each channel pair, similar to the HGB. Subsequently, we averaged the PLVs in a defined frequency range: the delta (0.1–3 Hz) or theta (3–7 Hz), in which we observed the most pronounced PLV activations (see Suppl. Fig. S8).

The ROI-level PLV (between *ROI_A_* and *ROI_B_*) was computed as a mean across all channel pairs of all subjects assigned to these two ROIs, yielding a 3-D matrix: *PLV*(*ROI_A_*, *ROI_B_*, *t*). Note that, to ensure a minimal degree of reproducibility, we included only those network connections with channel pairs from at least two subjects.

For both delta and theta PLV time curves (Fig. 7a), we selected three time points of interest (the onset, peak and offset) to illustrate the inter-regional PLV connectivity in Fig. 8 (but see also the Suppl. Movies V1 and V2 for the full temporal evolution of the PLVs). The peak is simply the local maximum of the PLV time course during the respective operand period (N1–N3). The onset was defined as the local maximum of the first derivative of the PLV; the offset as the local minimum. The derivative was estimated by three-point stencils, smoothed subsequently with a 5-Hz low-pass filter (Butterworth, 6th order, zero phase shift).

### Statistical testing for differences in distribution

Owing to heterogeneous and clinically determined electrode placement, classical subject-level group analyses were not adequate for all ROIs due to their relative undersampling (e.g., only four subjects with channels in IPS) and especially for some inter-regional interactions (see Suppl. Table T2). We therefore performed analyses at the channel level of each ROI, pooling channels assigned to the ROIs across patients and treated the trial-averaged channels (or channel pairs) as the unit of analysis, without modeling subject identity. This approach does not assume statistical independence across electrodes and does not support population-level inference across subjects; instead, results characterize response properties across the sampled intracranial recording sites.

For statistical testing, we used non-parametric tests, either the two-sided Wilcoxon rank-sum test or the two-sided, paired sign test, with FDR corrections for multiple testing^92^. More specifically, we used the paired sign test to assess the significance of difference between different conditions among either the subjects’ behavioral performance (Fig. S3) or the trial-averaged channels’ HGB activities (Figs. 5, S4, S6, S7). The Wilcoxon rank-sum test was used either to assess the significance of changes in the HGB activity against the baseline distribution (Fig. 4) across trial-averaged channels in each ROI, or to test for the significance of difference between selected ROIs (Figs. 6, 7, S5, S8, S9, S10). The use of the particular test is also reported in the figure legends.

## Data availability

The raw iEEG data that support the findings of this study are publicly available (https://zenodo.org/records/16778665). Further data are available from the corresponding authors upon reasonable request.

## Code availability

Code is publicly available for the iEEG data analysis (https://github.com/JiriHammer/SEEG_dataAnalysis).

## Acknowledgements

Both the Departments of Neurology and Pediatric Neurology, Second Faculty of Medicine, Charles University and Motol University Hospital are full members of ERN EpiCARE. All authors are members of the Epilepsy Research Centre Prague - EpiReC consortium. The authors would like to thank the patients for their participation in this study and CESNET for access to their data storage facility.

## Author contributions

MKa: Methodology, investigation, formal analysis, writing – original draft, visualization. AK: Investigation, data curation, writing – review & editing. BK: Writing – review & editing. VP: Investigation, writing – review & editing. JA: Writing - review & editing. RJ: Methodology, writing – review & editing. PJ: Data curation. DK: Investigation, data curation. MKu: Investigation, writing – review & editing. PK: Investigation, writing – review & editing. PM: Conceptualization, resources, writing – review & editing, supervision, funding acquisition. JH: Conceptualization, methodology, investigation, formal analysis, writing – original draft, visualization, supervision, funding acquisition.

