## Supplementary Material for "Temporal order of activations and interactions during arithmetic calculations measured by intracranial electrophysiological recordings in the human brain"

| Subject | Gender | Age | Handedness | Site of implantation ($N_{electrodes}/N_{contacts}$) | Epilepsy duration | Cognitive profile | Education | Task success rate (%) |
|---|---|---|---|---|---|---|---|---|
| P1 | M | 44 | R | L frontal precentral (6/66) | 36 | average | secondary | 88 |
| P2 | M | 36 | R | R temporo-parietal-insular (16/126) | 12 | above average | tertiary | 90 |
| P3 | F | 38 | R | bilateral fronto-insular (16/126) | 19 | average | tertiary | 78 |
| P4 | F | 28 | R | R fronto-temporal (11/126) | 10 | average | tertiary | 83 |
| P5 | M | 63 | R | L frontal + 2 electrodes in R frontal (15/192) | 46 | average | tertiary | 83 |
| P6 | F | 24 | L | R temporal (14/164) | 6 | average | primary | 65 |
| P7 | M | 49 | R | bilateral temporal + L frontal (11/126) | 27 | average | tertiary | 95 |
| P8 | F | 30 | L | bilateral temporal (14/186) | 19 | average | tertiary | 96 |
| P9 | M | 46 | R | R temporo-parieto-occipital (18/190) | 36 | below average | primary | 64 |
| P10 | M | 30 | L | R operculo-insular (15/189) | 28 | below average | secondary | 47 |
| P11 | F | 36 | R | L temporo-parieto-occipital (18/190) | 7 | average | tertiary | 88 |
| P12 | M | 36 | R | L temporo-insular (13/163) | 9 | below-average | secondary | 86 |
| P13 | M | 23 | L | L fronto-temporal (17/190) | 7 | average | secondary | 69 |
| P14 | F | 42 | R | R temporal (13/141) | 30 | average | secondary | 94 |
| P15 | F | 24 | R | L temporo-insular (13/159) | 1 | above average | tertiary | 90 |
| P16 | F | 23 | R | bilateral frontal mesial (14/178) | 7 | average | secondary | 92 |
| P17 | M | 51 | R | bilateral frontal (15/187) | 41 | average | secondary | 90 |
| P18 | M | 47 | R | R temporo-insular (14/167) | 13 | average | secondary | 90 |
| P19 | M | 18 | R | R temporo-parietal-insular (15/149) | 4 | average | primary | 85 |
| P20 | M | 36 | R | R temporo-parietal (13/125) | 29 | below average | secondary | 75 |

**Supplementary Table T1: Subject details.** Twenty epilepsy-surgery candidates (P1–P20) participated in the task. The columns represent: demographic information; handedness; implantation scheme, specifying the implanted hemisphere (right/left/bilateral implantation) and the number of implanted electrodes and number of recording contacts (in parentheses), epilepsy duration; epileptogenic pathology, if known from histology or suspected based on neuroimaging; cognitive profile, based on presurgical neuropsychological assessment using an abridged battery of subtests from the WAIS-III (vocabulary, similarities, block-design, matrix reasoning ); level of education; and success rate in the sequential arithmetic task. Abbreviations: F, female; M, male; R, right; L, left.

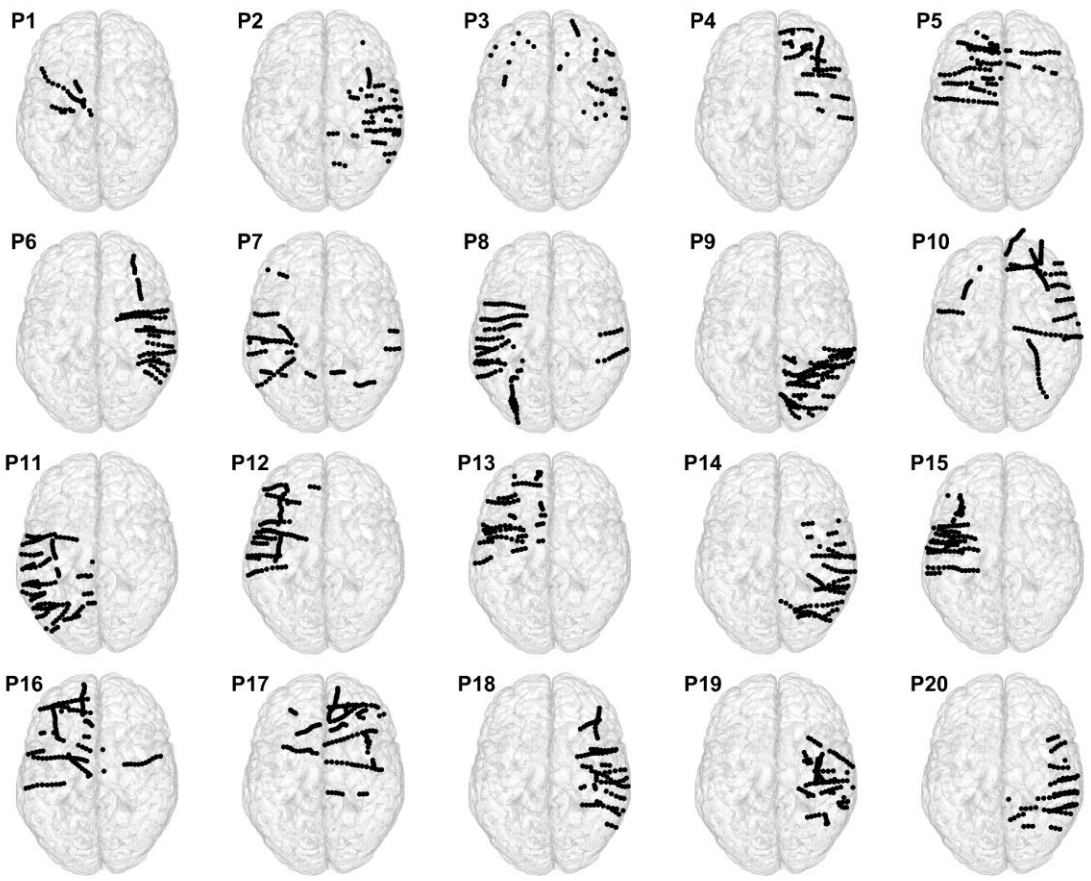


**Supplementary Figure S1: Implantation schemes of the subjects P1–P20.** Circles represent channels projected on a brain template (Colin27, top view). The channel location corresponds to the centre of the MNI coordinates of the bipolar-referenced SEEG electrode contact pair. Note that due to the normalization, the trajectory of channels may appear slightly bent; in reality, they were straight.

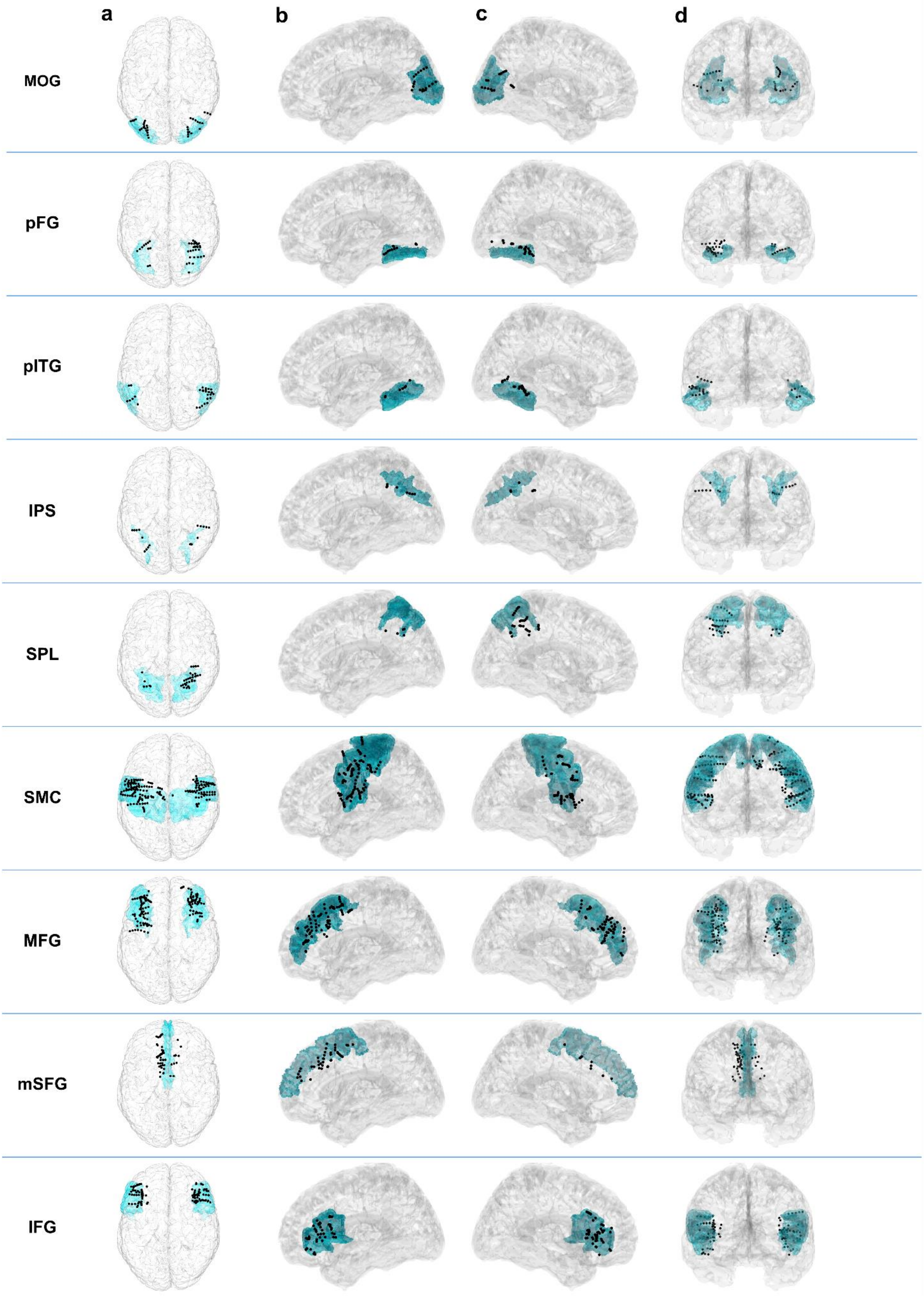

a
b
c
d
MOG
pFG
pITG
IPS
SPL
SMC
MFG
mSFG
IFG

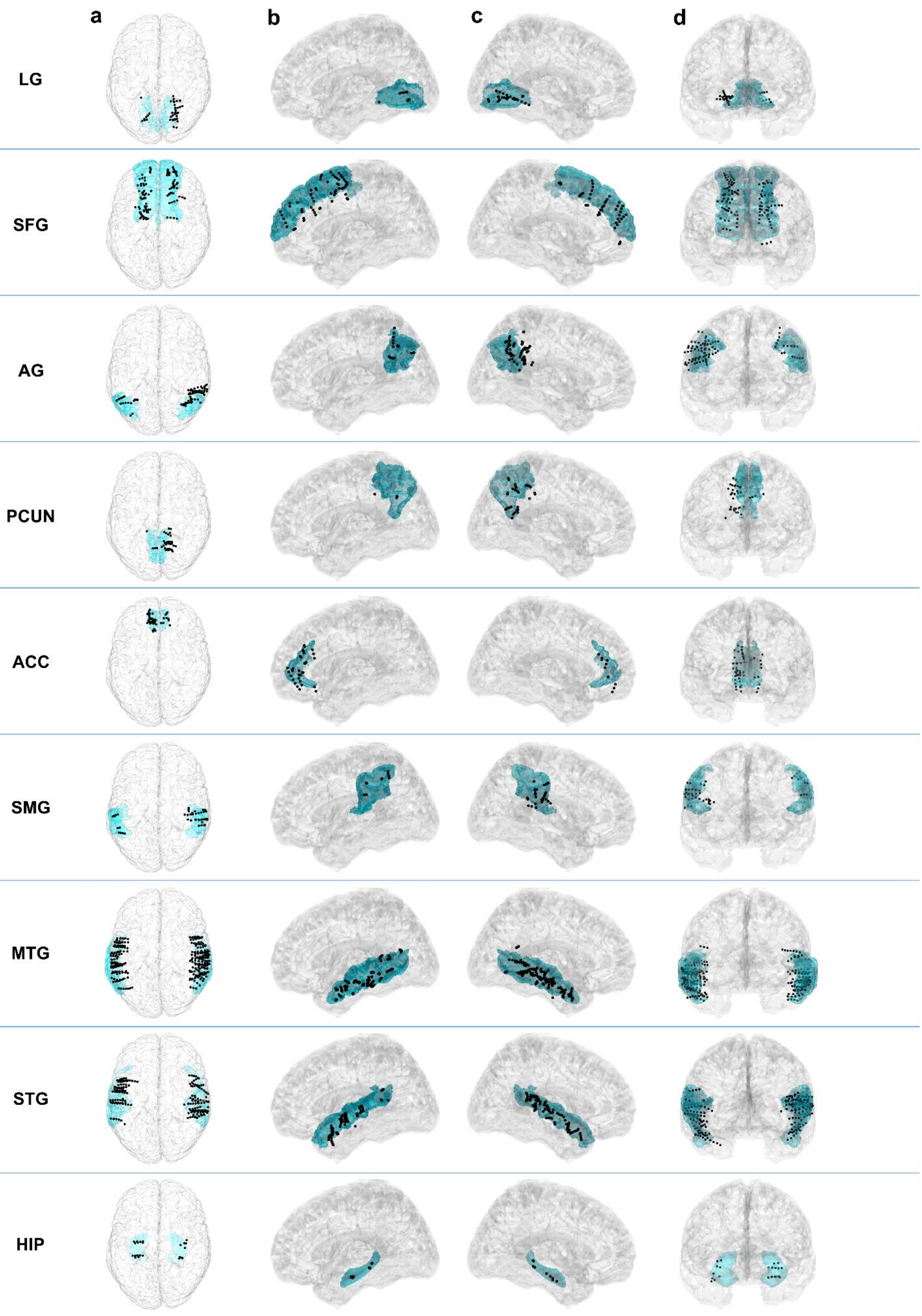


**Supplementary Figure S2: Channel localisation in the arithmetic network ROIs.** All channels assigned to a particular ROI from all subjects are depicted as black dots, the ROI is highlighted in blue, its name indicated on the left. The position of the channels (i.e., centre of the bipolar pairs) is shown as projections on a transparent brain template (Colin27) in **(a)** top view, **(b)** left lateral, **(c)** right lateral, **(d)** coronal view (either from front or back, depending on the ROI to ensure optimal visibility). Note that the anatomical assignment of the channels to each ROI was done on individual, real brain scans; due to the normalization to the brain template, projections and the channel inclusion criteria, where we required that at least one of the bipolar contacts was localized in the ROI, some channels may appear slightly out of the ROI.

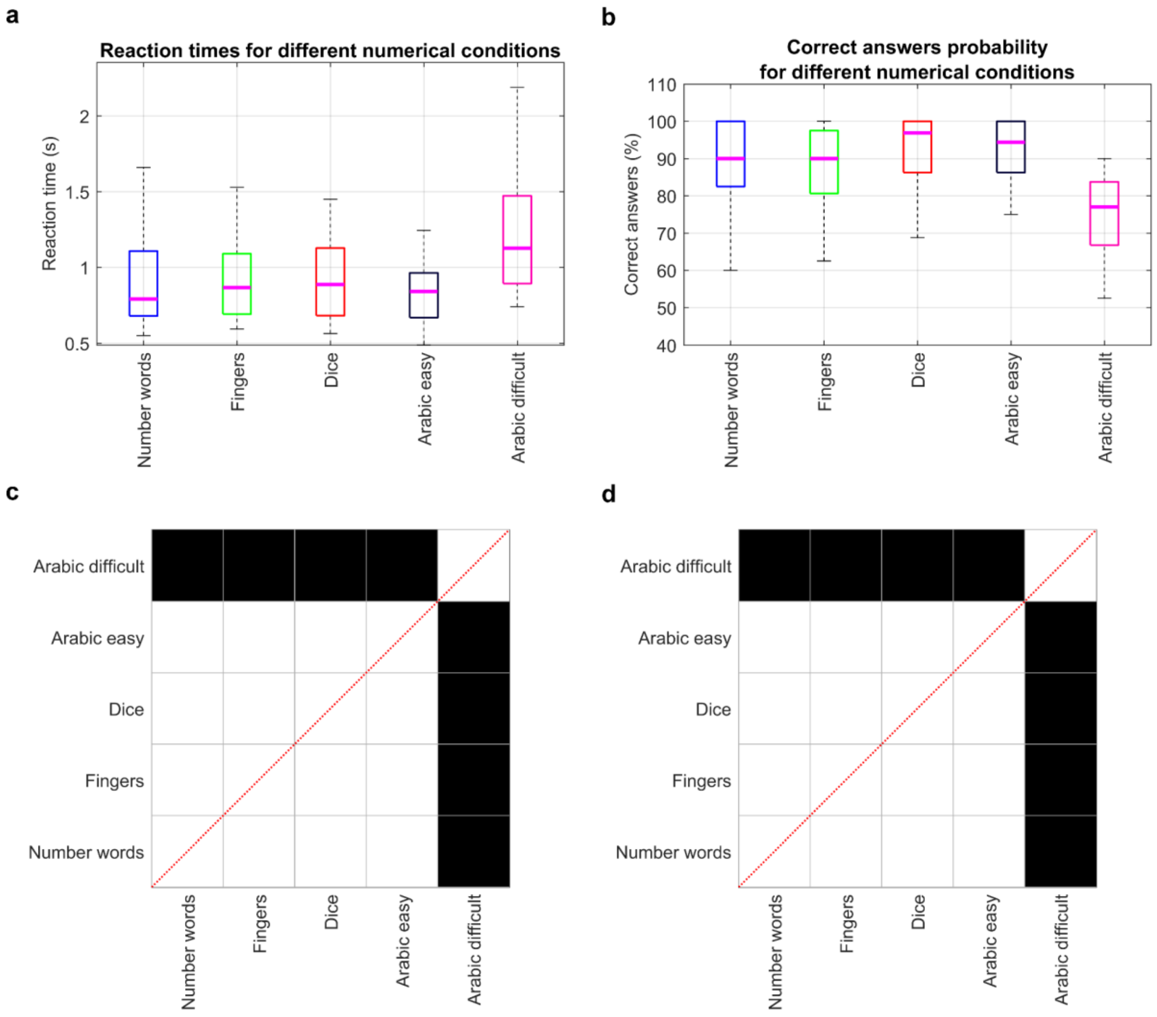


**Supplementary Figure S3: Behavioral responses on task conditions.** **(a)** Reaction times and **(b)** ratio of correct answers for each task condition differing in task difficulty and presented numerical format: number words, fingers, dice, Arabic easy and Arabic difficult (see Methods for more details). Statistical comparison of differences in the distribution of subject means for **(c)** reaction times or **(d)** correct answers ratio among all conditions (sign test, black: $P < 0.05$, FDR corrected). As expected, the subjects were slower and made more errors during the more demanding condition.

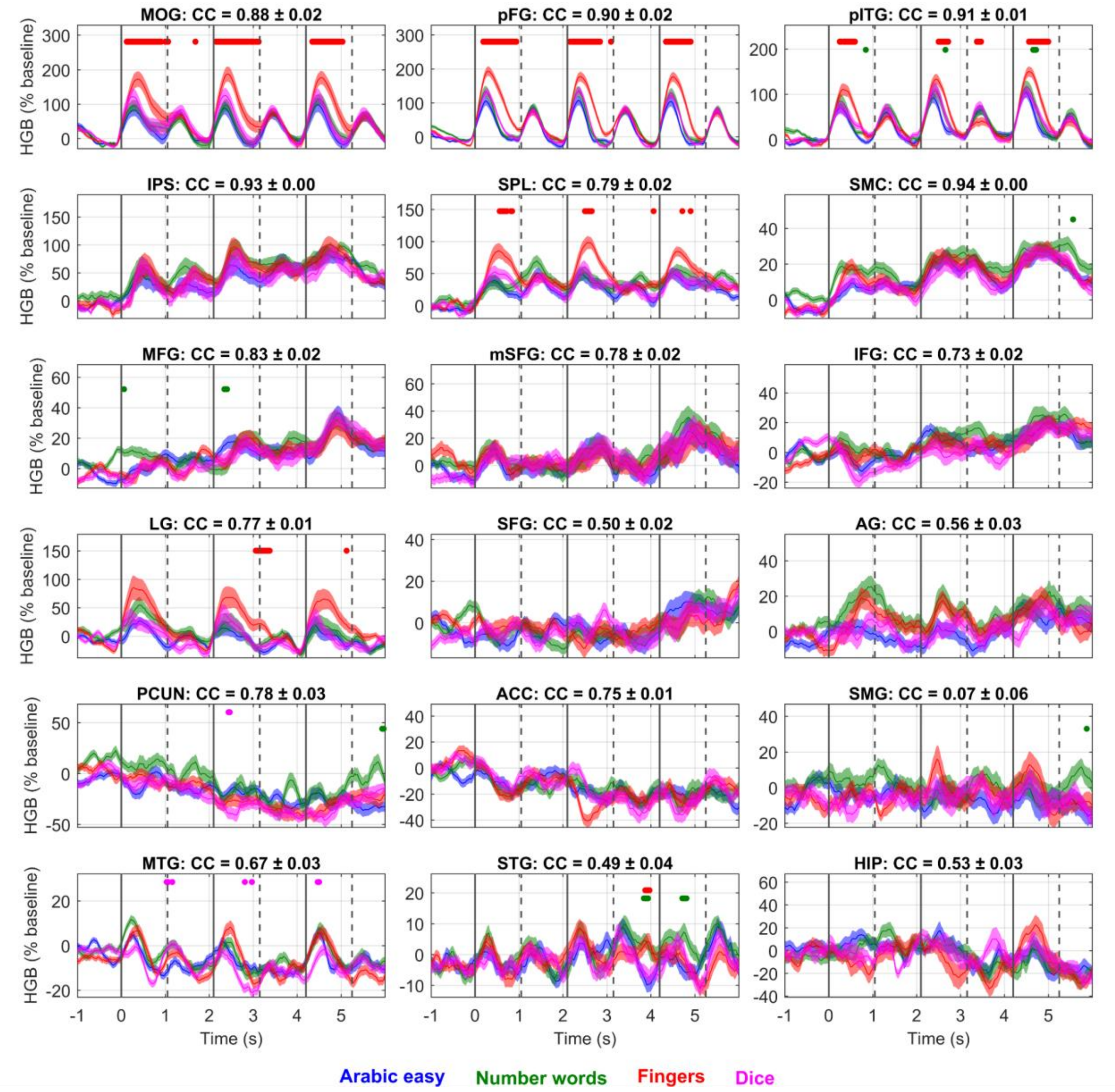


**Supplementary Figure S4: HGB activity for different numerical formats.** To assess the format-dependence of the HGB responses, the operands in the easier equations (result < 10) were presented in different numerical formats: Arabic numbers (blue), number words (green), fingers (red) and dice (magenta). Each subplot represents one ROI (same conventions as in Fig. 1) with time-resolved (x-axis) HGB activities (y-axis). We compared the HGB corresponding to Arabic numbers against all other numerical formats using the paired, two-sided sign test to assess the numerical format-related differences across the trial-averaged channels in each ROI. A colored dot above indicates the significance of difference ($P < 0.001$, FDR corrected) between the Arabic numbers and the numerical format with corresponding color (e.g., a red dot indicates that HGB responses of fingers were significantly different from the HGB responses during the presentation of the Arabic numbers). For each ROI, we computed the mean (± SEM) correlation coefficients (CC) among all pairs of the numerical formats, the value indicated in the title above each subplot. Fingers elicited significantly higher HGB activations in the VOTC, which could be attributed to their higher stimulus complexity. The HGB activations in the IPS appeared to be largely format-independent. Overall, the HGB activity was highly correlated for the different numerical formats.

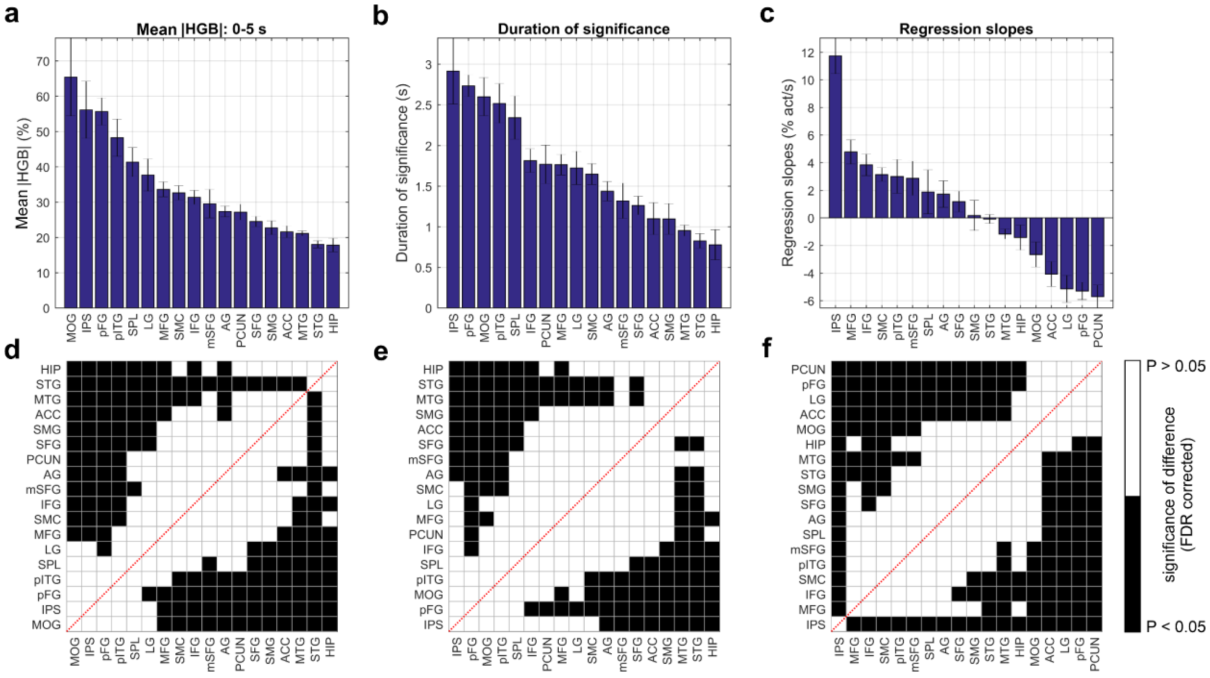


**Supplementary Figure S5: Comparison of HGB activity indicators among the ROIs of the arithmetic network.** We selected three different indicators for the single-channel, trial-averaged HGB activity in the time interval between 0 and 5 s after the presentation of the first operand: **(a)** mean (± SEM) of |HGB|, **(b)** duration of significance, and **(c)** slope of the linear regression of the HGB activity. Bottom row **(d-f)**: significance of differences between all ROI pairs presented on a chessboard plot (black: $P < 0.05$, Wilcoxon rank-sum test, FDR corrected; else white) for each indicator. The indicators confirm the IPS, SPL, pITG, pFG, MOG, several frontal regions (MFG, SFGM, and IFG), and SMC as the key areas of the arithmetic network.

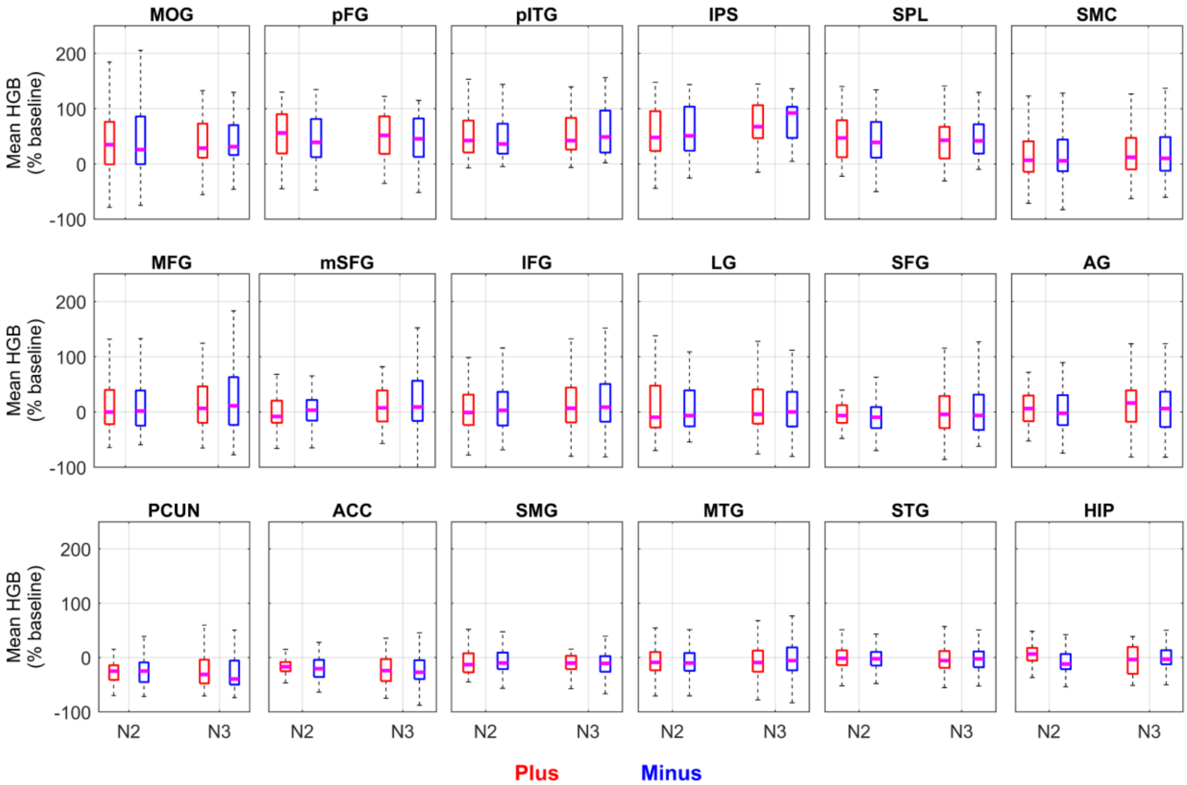


**Supplementary Figure S6: Mean HGB responses in the arithmetic network following plus or minus operators.** Each subplot corresponds to the selected ROI in the arithmetic network indicated by the title above (same notations as in Fig. 1). The mean HGB responses were calculated as the mean across each operand period from the time-resolved, trial-averaged HGB activity across channels in each ROI. We compared the mean HGB responses (y-axis) during the operand periods (x-axis: either N2 or N3) following the plus (red boxplots) or minus (blue boxplots) operators. We tested the HGB values across the trial-averaged channels (plus vs. minus) at both operand periods (paired, two-sided sign test; P-level = 0.05; FDR corrected for multiple testing), but found no statistically significant difference in the HGB responses.

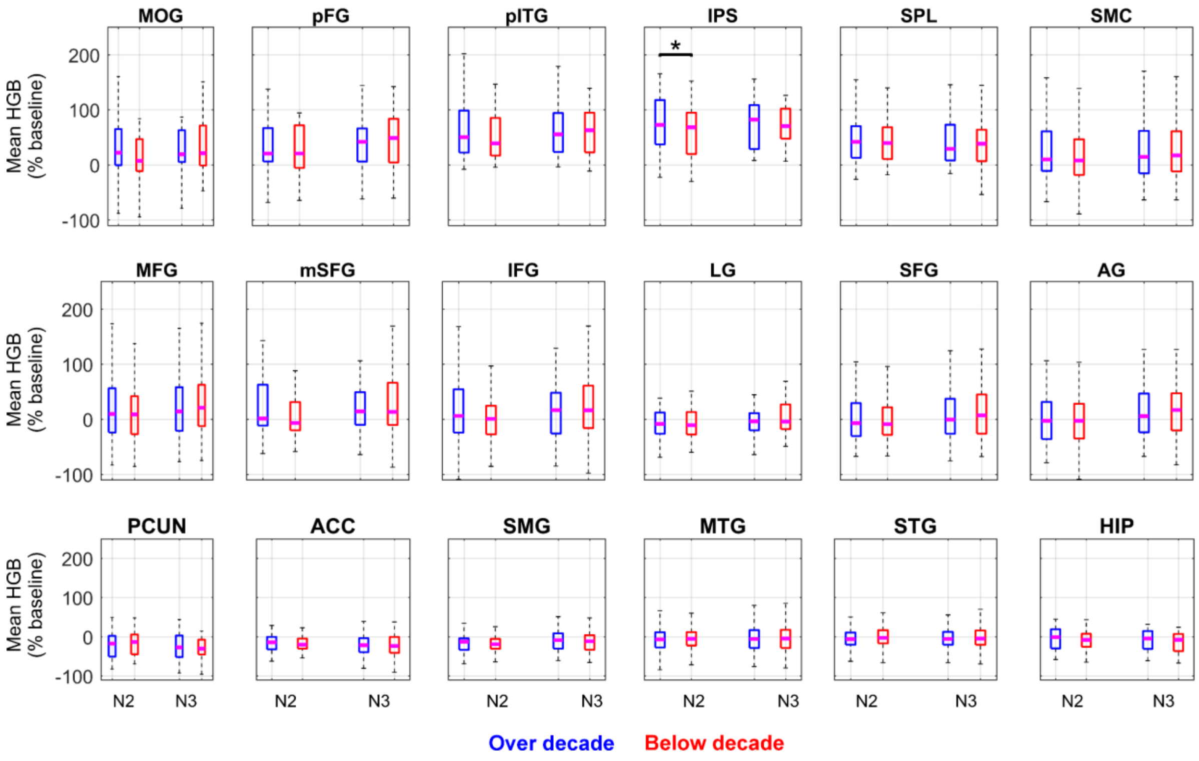


**Supplementary Figure S7: Cross-decade HGB responses in the arithmetic network.** During the task, subjects also solved more difficult arithmetic equations (condition: Arabic difficult), where the intermediate result crossed the decade exactly once in each equation. The cross-decade calculation occurred with equal probability at both operands N2 and N3. Blue boxplots correspond to the condition when the intermediate result was over the decade (e.g., 45+6 or 45-6), while red boxplots correspond to the condition when the intermediate result was not over the decade (e.g., 45+4 or 45-4). The analysis was split between the cross-decade responses following either the second (N2) or the third (N3) operators separately, due to cumulative activity in some regions across the entire equation (see Fig. 4: the IPS for example). Subplots represent each ROI with a title indicating its name (same conventions as in Fig. 1). We tested for significance of difference between the cross-decade conditions (i.e., blue against red) using the paired, two-sided sign test across the trial-averaged channels. The only ROI with a significant cross-decade effect was the IPS ($P < 0.05$, FDR corrected).

| | MOG | pFG | LG | pITG | MTG | STG | IPS | SMG | AG | PCUN | SPL | ACC | SMC | HIP | SFG | mSFG | MFG | IFG |
|---|---|---|---|---|---|---|---|---|---|---|---|---|---|---|---|---|---|---|
| IFG | 86 (2) | 42 (2) | | 38 (3) | 1060 (10) | 537 (8) | 68 (2) | 150 (5) | 125 (4) | 21 (3) | 24 (3) | 397 (7) | 1112 (11) | 138 (6) | 734 (7) | 344 (5) | 1004 (12) | |
| MFG | 84 (2) | | | 15 (2) | 1046 (9) | 562 (7) | | 159 (4) | 137 (3) | 59 (2) | 86 (2) | 706 (7) | 2110 (12) | 188 (5) | 1823 (8) | 972 (6) | | 1004 (12) |
| mSFG | | | | | 99 (2) | | | | | | | 386 (3) | 893 (4) | 41 (2) | 844 (6) | | 972 (6) | 344 (5) |
| SFG | | | | | 127 (4) | 45 (2) | | 68 (2) | | | | 774 (6) | 1882 (7) | 21 (3) | | 844 (6) | 1823 (8) | 734 (7) |
| HIP | | 53 (3) | | 22 (2) | 312 (8) | 292 (7) | | 38 (2) | 62 (4) | 9 (2) | | 15 (3) | 236 (4) | | 21 (3) | 41 (2) | 188 (5) | 138 (6) |
| SMC | 170 (3) | 57 (3) | 239 (3) | 100 (4) | 1579 (10) | 713 (8) | 172 (2) | 338 (5) | 352 (5) | 120 (4) | 123 (3) | 1061 (6) | | 236 (4) | 1882 (7) | 893 (4) | 2110 (12) | 1112 (11) |
| ACC | | | | | 139 (5) | 89 (3) | | 53 (2) | | | | | 1061 (6) | 15 (3) | 774 (6) | 386 (3) | 706 (7) | 397 (7) |
| SPL | 113 (2) | 76 (3) | 269 (2) | 29 (3) | 580 (4) | 330 (4) | 49 (2) | 68 (4) | 614 (4) | 434 (5) | | | 123 (3) | | | | 86 (2) | 24 (3) |
| PCUN | 100 (3) | 119 (5) | 239 (4) | 91 (5) | 565 (6) | 347 (6) | 59 (3) | 113 (5) | 593 (6) | | 434 (5) | | 120 (4) | 9 (2) | | | 59 (2) | 21 (3) |
| AG | 304 (5) | 262 (7) | 495 (6) | 173 (5) | 1478 (10) | 966 (9) | 160 (4) | 333 (7) | | 593 (6) | 614 (4) | | 352 (5) | 62 (4) | | | 137 (3) | 125 (4) |
| SMG | 82 (4) | 112 (6) | 177 (4) | 106 (5) | 632 (8) | 378 (6) | 80 (4) | | 333 (7) | 113 (5) | 68 (4) | 53 (2) | 338 (5) | 38 (2) | 68 (2) | | 159 (4) | 150 (5) |
| IPS | 84 (3) | 39 (3) | 115 (3) | 36 (2) | 303 (4) | 97 (3) | | 80 (4) | 160 (4) | 59 (3) | 49 (2) | | 172 (2) | | | | | 68 (2) |
| STG | 447 (5) | 526 (8) | 324 (5) | 329 (6) | 3464 (14) | | 97 (3) | 378 (6) | 966 (9) | 347 (6) | 330 (4) | 89 (3) | 713 (8) | 292 (7) | 45 (2) | | 562 (7) | 537 (8) |
| MTG | 1031 (6) | 758 (8) | 644 (6) | 595 (6) | | 3464 (14) | 303 (4) | 632 (8) | 1478 (10) | 565 (6) | 580 (4) | 139 (5) | 1579 (10) | 312 (8) | 127 (4) | 99 (2) | 1046 (9) | 1060 (10) |
| pITG | 110 (3) | 191 (5) | 147 (3) | | 595 (6) | 329 (6) | 36 (2) | 106 (5) | 173 (5) | 91 (5) | 29 (3) | | 100 (4) | 22 (2) | | | 15 (2) | 38 (3) |
| LG | 180 (5) | 119 (5) | | 147 (3) | 644 (6) | 324 (5) | 115 (3) | 177 (4) | 495 (6) | 239 (4) | 269 (2) | | 239 (3) | | | | | |
| pFG | 143 (5) | | 119 (5) | 191 (5) | 758 (8) | 526 (8) | 39 (3) | 112 (6) | 262 (7) | 119 (5) | 76 (3) | | 57 (3) | 53 (3) | | | | 42 (2) |
| MOG | | 143 (5) | 180 (5) | 110 (3) | 1031 (6) | 447 (5) | 84 (3) | 82 (4) | 304 (5) | 100 (3) | 113 (2) | | 170 (3) | | | | 84 (2) | 86 (2) |

**Supplementary Table T2: Number of channel pairs and subjects among the ROIs of the arithmetic network used in the PLV connectivity analysis.** The text in each cell of the colour-coded table denotes the number of the iEEG channel pairs (upper number) and the number of subjects (lower number in brackets). Note that for reproducibility of the results, we demanded that at least two subjects contribute to any given connections. Hence, some ROI pairs were not analyzed due to undersampling, and are represented by empty cells.

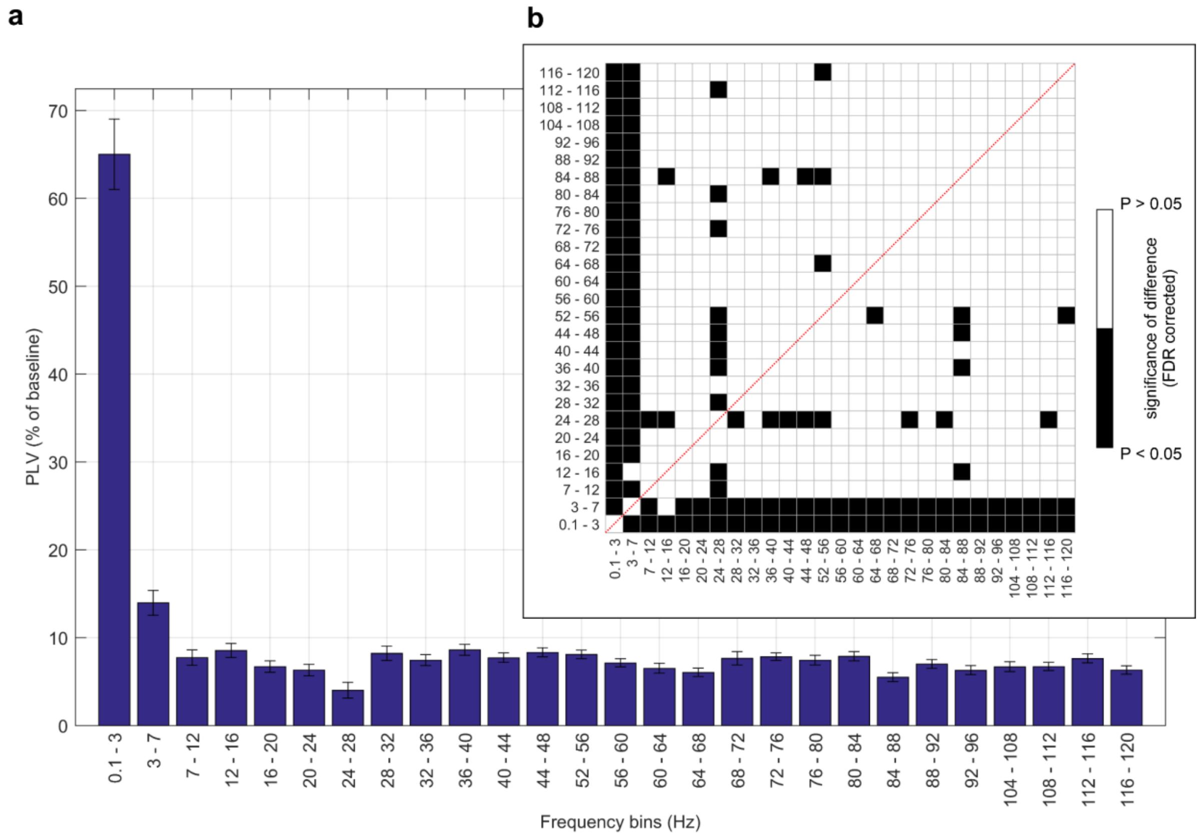


**Supplementary Figure S8: Grand average of PLVs for different frequency bands.** The PLVs were computed in different frequency bands using the filter-Hilbert approach. The grand average represents the sum of the PLVs across all 137 ROI pairs and the entire time period from 0–6 s after the presentation of the first operand, N1. **(a)** For each frequency bin (x-axis), we computed the grand average of the PLVs. Each bar represents PLVs in one frequency bin (mean ± SEM across all 137 ROI pairs included into the analysis, provided that at least two subjects contributed to the ROI pair). The PLV of each ROI-pair was, in turn, computed as a mean across all respective channel pairs. We divided the frequency spectrum of the iEEG signal into 4-Hz bins, with the exception of low frequencies, for which the binning followed the traditional division into delta (0–3 Hz), theta (3–7 Hz), and alpha (7–12 Hz) bands. **(b)** Statistical comparison of PLVs by frequency bin, with significant differences marked in black ($P < 0.05$, Wilcoxon rank sum test, FDR corrected). By far, the strongest PLV effects were observed in the delta band, followed by the theta band.

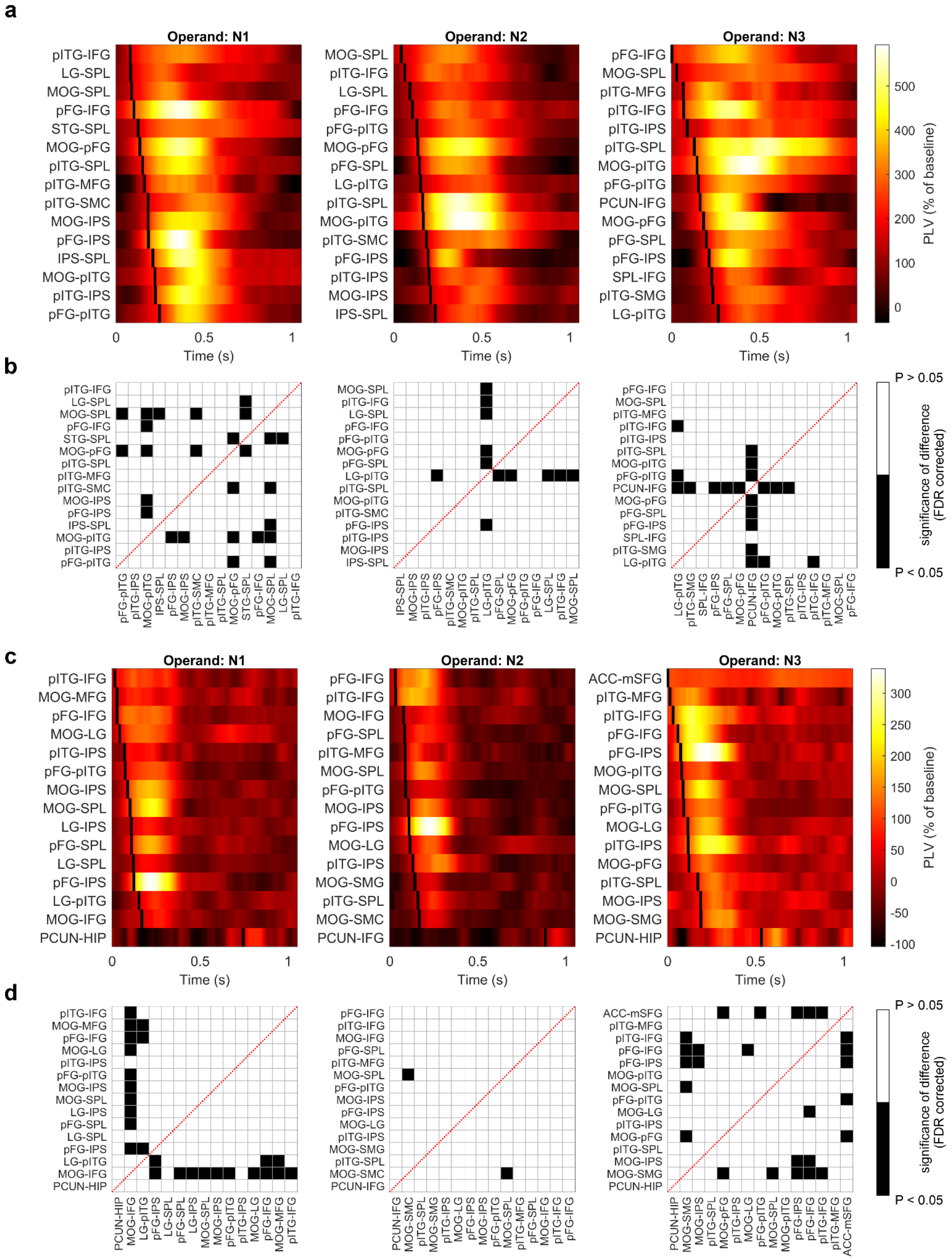


**Supplementary Figure S9: Temporal order of the delta (0.3–3 Hz) and theta (3–7 Hz) PLV onsets within the arithmetic network.** As in Fig. 7, we selected 15 pairs of ROIs in the arithmetic network based on their maximum PLV value for each operand period (N1–N3; figure columns) and ordered them according to the latencies of the onset. For a given ROI pair, the PLV onset was defined as the time point when the PLV time-curve crossed 50% of its peak value in each operand period, provided that the peak value was above 3$\sigma$ of the baseline PLV distribution. **(a)** Temporal profiles (x-axis) of the delta PLVs between the 15 strongest ROI pairs (rows of the subplots; names indicated on the y-axis). The PLVs were colour-coded. **(b)** The chessboard plots the significance of the maxima latency differences between all ROI pairs (black: P < 0.05, Wilcoxon rank-sum test, FDR corrected; else white). **(c, d)** Same as in a, b, but for the theta band. Surprisingly, the earliest PLV connections were found between VOTC and frontal areas (pITG–IFG, pFG–IFG, pITG–MFG).

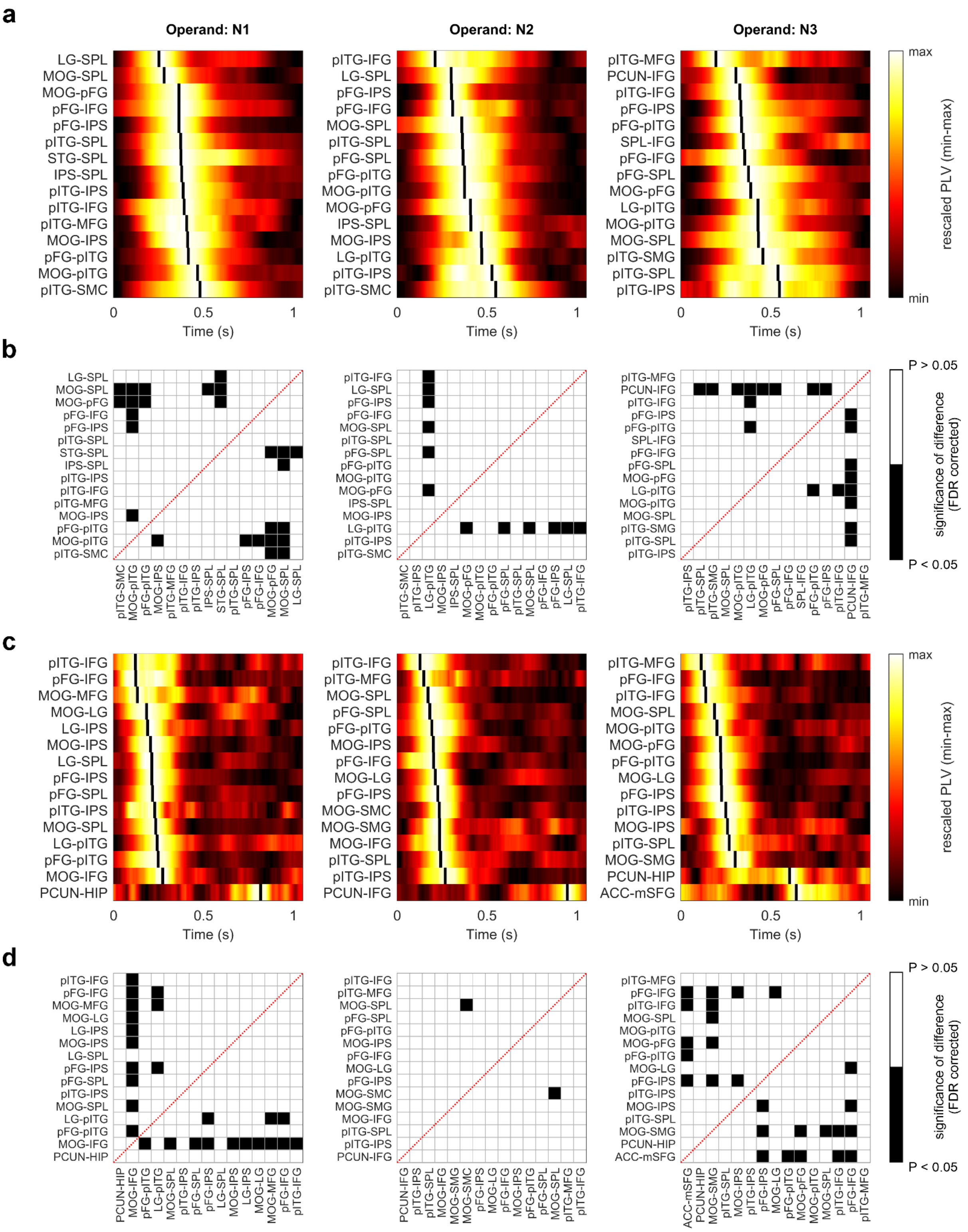

**Supplementary Figure 10: Temporal order of the delta (0.3–3 Hz) and theta (3–7 Hz) PLV peaks within the arithmetic network.** As in Fig. 7, 15 pairs of ROIs in the arithmetic network based on their maximum PLV value for each operand period (N1–N3; figure columns) were selected and ordered according to the latencies of their peaks (i.e., local maxima in each operand period), provided that the peak value was above 3$\sigma$ of the baseline PLV distribution. **(a)** Temporal profiles (x-axis) of the delta PLVs, same as in Fig. 7b, but here rescaled to the minimum and maximum of each operand period for a clearer visualization/comparison of the PLV temporal profiles of the selected ROI-pairs (rows of the subplots; names indicated on the y-axis). **(b)** The chessboard plots the significance of the delta PLV maxima latency differences between all ROI pairs (black: $P < 0.05$, Wilcoxon rank-sum test, FDR corrected; else white). **(c, d)** Same as in a, b, but for the theta band. Again, the earliest PLV peaks were found between VOTC and frontal areas (pITG–IFG, pFG–IFG, pITG–MFG).

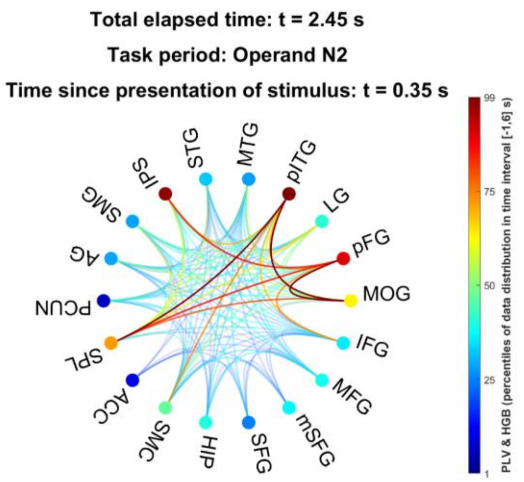


**Supplementary Movie V1: Animation of the HGB and FC changes in the sequential arithmetic task: delta band (0.1–3 Hz) PLV.** Each dot represents an ROI (or node) of the arithmetic network, color-coded based on its HGB activity at a given time, *t*. Each line represents the connection (or edge) between the ROIs, colour-coded based on the delta PLV activity at the time *t*. The colour-scale is common to both HGB and PLV, and spans the 1–99th percentile of the respective data distribution. The text above the figure shows the total elapsed time, where $t = 0$ s corresponds to the presentation of the first operand, the specification of the task period (baseline, operand N1–N3, or operator O1-O3), and time since the presentation of the stimulus (operand or operator).

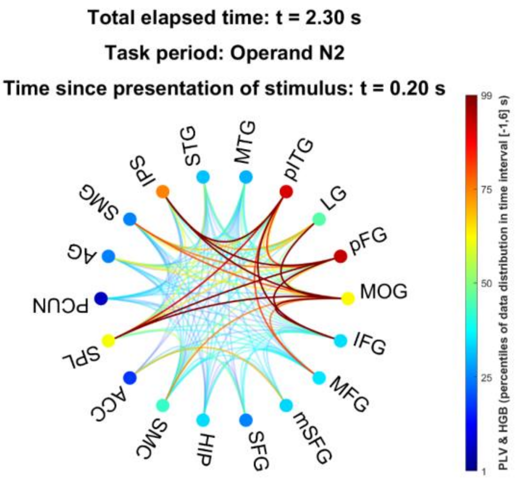


**Supplementary Movie V2: Animation of the HGB and FC changes in the sequential arithmetic task: theta band (3–7 Hz) PLV.** Each dot represents an ROI (or node) of the arithmetic network, color-coded based on its HGB activity at a given time, *t*. Each line represents the connection (or edge) between the ROIs, colour-coded based on the theta PLV activity at the time *t*. The colour-scale is common to both HGB and PLV, and spans the 1–99th percentile of the respective data distribution. The text above the figure shows the total elapsed time, where *t* = 0 s corresponds to the presentation of the first operand, the specification of the task period (baseline, operand N1–N3, or operator O1-O3), and time since the presentation of the stimulus (operand or operator).

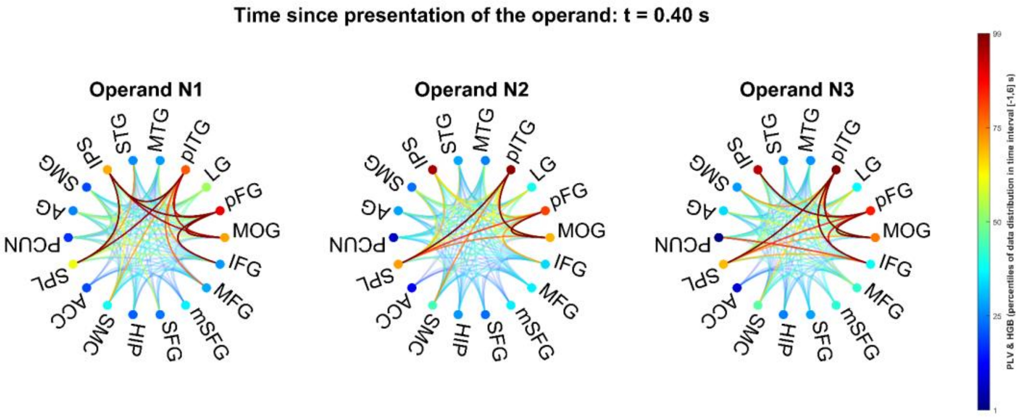


**Supplementary Movie V3: Animation of the HGB and FC changes for the three operand periods: delta band (0.1–3 Hz) PLV.** Each subplot shows the HGB and delta PLV changes after the presentation of the operand N1–N3 (indicated in the title above). In each subplot, a dot represents an ROI (or node) of the arithmetic network, colour-coded based on its HGB activity at a given time, *t*. Each line represents the connection (or edge) between the ROIs, colour-coded based on their PLV activity at the time *t*. The colour-scale is common to both HGB and PLV, and spans the 1–99th percentile of the respective data distribution. The text above the figure shows the elapsed time since the presentation of each operand.

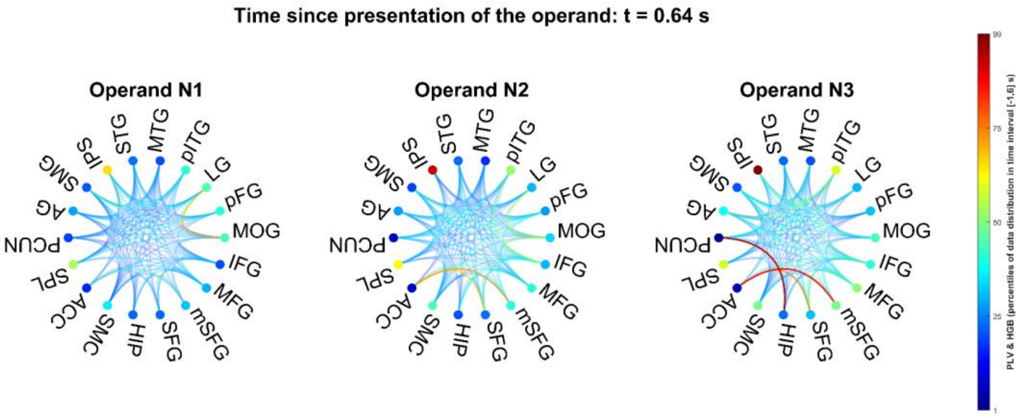


**Supplementary Movie V4: Animation of the HGB and FC changes for the three operand periods: theta band (0.1–3 Hz) PLV.** Each subplot shows the HGB and theta PLV changes after the presentation of the operand N1–N3 (indicated in the title above). In each subplot, a dot represents an ROI (or node) of the arithmetic network, colour-coded based on its HGB activity at a given time, *t*. Each line represents the connection (or edge) between the ROIs, colour-coded based on their PLV activity at the time *t*. The colour-scale is common to both HGB and PLV, and spans the 1–99th percentile of the respective data distribution. The text above the figure shows the elapsed time since the presentation of each operand.
